# Genomic characterization of type I-E* CRISPR-Cas systems in *Klebsiella pneumoniae*: association of spacer diversity and antimicrobial resistance

**DOI:** 10.64898/2025.12.03.692015

**Authors:** Ankur Rao, Tawsif Ahmed Kazi, Sharmi Naha, Dipanjan Roy, Suchandra Mukherjee, Dipanjan Ghosh, Sulagna Basu

## Abstract

**Background:** *Klebsiella pneumoniae,* known to acquire antimicrobial resistance genes (ARGs) through horizontal gene transfer, possess type I CRISPR-Cas systems. This study investigated the association of multidrug resistance (MDR) and CRISPR-Cas in septicemic *K. pneumoniae*, along with comparative analysis of CRISPR-positive global *K. pneumoniae*.

**Methods:** Archived *K. pneumoniae* from septicemic neonates (2008-2020) were assessed for CRISPR-Cas system, *cas* gene expression and correlation with ARG carriage. Whole-genome sequencing was performed and diversity of spacers, protospacers adjacent motifs (PAMs), targeted plasmids, were evaluated. Comparative analyses were performed for CRISPR-positive global *K. pneumoniae*.

**Results:** Twenty-five percent of study *K. pneumoniae* possessed complete type I-E* CRISPR-Cas comparable to 22% of global *K. pneumoniae*, which were distributed equally to type I-E & I-E*. CRISPR-positive study strains (83%) were MDR, and exhibited no significant association of CRISPR-Cas with MDR phenotype. Spacers noted in non-MDRs were diverse, showed increased average count and identified more diverse PAMs compared to spacers from MDR strains. Plasmids harboring ARGs such as *bla*_NDM_, *bla*_CTX-M_, *qnrS1*, *mcr-8*, etc. were frequently targeted by spacers from non-MDRs. Presence of CRISPR-Cas system in epidemic MDR clones such as ST14/15/23 in I-E* (local, global) and ST147 in I-E (global), indicated a lineage-specific association with CRISPR-Cas subtypes.

**Conclusion:** This study demonstrated the co-existence of CRISPR-Cas systems within both MDR and non-MDR *K. pneumoniae*, with no significant correlation with either group. However, presence of diverse spacers in non-MDR strains that identified more PAMs and targeted ARG-bearing plasmid implied role of CRISPR-Cas system in limiting ARG acquisition.

## 1. Introduction

*Klebsiella pneumoniae* is a major nosocomial pathogen responsible for pneumonia, urinary tract, wound, cystitis, sepsis and bloodstream infections (1). World Health Organization (WHO) has included third-generation cephalosporin- and carbapenem-resistant *K. pneumoniae,* as a critical pathogen (2) in the priority pathogen list, underscoring the urgent need to control their spread and mitigate antimicrobial resistance (AMR).

The emergence of resistance to multiple antimicrobials, including carbapenems-regarded as antibiotics of ‘last resort’- has been primarily driven by the horizontal transfer (HGT) of AMR genes encoded on plasmids (3). HGT facilitates the dissemination of these resistance determinants across diverse bacterial species, enabling survival under antibiotic selection pressure. However, the acquisition of exogenous DNA imposes a fitness cost on recipient bacteria (4). To counterbalance this cost, bacteria have evolved sophisticated defense systems, notably the Clustered Regularly Interspaced Short Palindromic Repeats (CRISPR) and associated Cas (CRISPR-associated) proteins. This adaptive immune system provides protection against invading genetic elements such as bacteriophages and plasmids, including those harboring AMR genes, thereby modulating gene flow within microbial populations (5, 6).

The CRISPR-Cas system consists of three main components: the CRISPR array, *cas* genes and a leader sequence. The CRISPR array contains short, repeating DNA sequences separated by spacers derived from foreign genetic material, allowing bacteria to recognize and restrict subsequent invasions of the same genetic elements (7). The CRISPR-Cas system works in three phases: adaptation, expression and interference. In the adaptation phase, DNA from invaders such as bacteriophages or plasmids is integrated into the CRISPR array as spacers. During expression, the array is transcribed into CRISPR RNAs (crRNAs), which direct Cas proteins to target and cleave foreign DNA during the interference phase (8, 9). The spacer sequences, ranging from 21 to 47 bp in size (10) present between the direct repeat sequences are responsible for specific sequence-based targeting of foreign DNA sequences. This function is useful for targeting antimicrobial resistance genes (ARGs) harbored on foreign elements such as plasmids.

On the basis of *cas* subunit complexity, CRISPR is divided into two classes, Class 1 (multi-subunit complex) and Class 2 (single subunit complex). Further classification into types exists on the basis of the signature *cas* gene– currently, six types have been reported (I-VI); of which types I, III and IV belong to Class 1 and types II, V and VI belong to Class 2 (11). Complete CRISPR-Cas systems are those which possess both *cas* genes as well as the spacer sequences (12). In bacteria, type I is the most common CRISPR-Cas system identified which comprises of both *cas1* and *cas3* genes responsible of spacer acquisition and interference (13). *K. pneumoniae* in particular, is known to possess type I-E and I-E* systems with variation in the arrangement (14). The differences between these two subtypes are in the arrangement of the three *cas* genes (*cas5*, *cas6* and *cas7*). In I-E subtype, these 3 *cas* genes are arranged as *cas7-cas5-cas6*, while in I-E*, they are arranged as *cas6-cas7-cas5* (12).

The relationship between CRISPR and AMR is very complex and often species specific (13–16). In the last decade, most studies on *K. pneumoniae* have reported an inverse correlation between presence of CRISPR-Cas systems and antimicrobial resistance (14–19). However, contradicting results have been reported in other organisms such as *Escherichia coli* (20) and *Campylobacter jejuni* (21). Given this variability, more research is needed to resolve this paradox and better understand the potential role of CRISPR-Cas systems in controlling the spread of ARGs. Such insights are essential for developing innovative strategies to combat AMR. This study aims to investigate the CRISPR-Cas systems in *K. pneumoniae* strains, the acquired spacers and their targets and analyze their functional role in relation to AMR.

## 2. Materials and Methods

### 2.1. Identification and antimicrobial susceptibility testing (AST) of study strains

Archived *K. pneumoniae* (N=241) isolated from blood of septicemic neonates collected during 2008 to 2020 were identified using biochemical tests and VITEK^®^ 2 Compact system (bioMérieux, Marcy-l’Étoile, France). Antimicrobial susceptibility tests were performed by Kirby-Bauer disk-diffusion method for strains from 2008-2017, while for strains from 2018 to 2020, VITEK^®^ 2 Compact system AST cards were used. Results were analyzed according to Clinical & Laboratory Standards Institute (CLSI), 2025 (22) for all antimicrobials, except tigecycline, which was interpreted in accordance with European Committee on Antimicrobial Susceptibility Testing (EUCAST) guidelines, 2025 (23). Strains resistant to three or more antimicrobial groups were defined as multidrug-resistant (MDR), and those resistant to less than three antimicrobial groups were non-MDR (24).

### 2.2. Genotypic characterization of study strains

Different antimicrobial resistance genes– extended spectrum β-lactamase genes (ESBLs), carbapenemases, plasmid-mediated AmpC genes (pAmpC), aminoglycoside and fluoroquinolone resistance genes were detected by polymerase chain reaction (PCR) as mentioned previously (25). These genes were further Sanger sequenced as described previously (25) using Applied Biosystems ProFlex 3 x 32-well PCR system (Thermo Fisher Scientific, United States).

### 2.3. Identification and characterisation of CRISPR-Cas systems

*cas1* and *cas3* genes responsible for spacer acquisition and excision of exogenous DNA sequences present in type I CRISPR-Cas systems were amplified using Applied Biosystems Proflex 3 x 32-well PCR System (Thermo Fisher Scientific, USA). Primer for *cas1* was designed in this study (Table S1) and primer sequence for *cas3* primer was adopted from a previous study (18).

### 2.4. Pulsed-Field Gel Electrophoresis (PFGE) for detection of distinct strains

For all complete CRISPR-Cas positive (with both *cas1* and *cas3* present) strains, PFGE was performed as a prerequisite to prevent inclusion of clonal strains. PFGE was performed following standardised PulseNet procedure (http://www.cdc.gov/pulsenet/protocols.html). Distinct strains were determined by visual detection of band patterns using the Tenover criterion (26).

### 2.5. Real time quantitative PCR (RT-qPCR) analysis of *cas* genes in MDR and non-MDR strains

For a type I CRISPR-Cas system to be functional, *cas1* and *cas3* needs to be active. Hence, understanding the expression of these two genes is important and gene expression analysis of the same was performed using RT-qPCR. Expression of *cas1* and *cas3* genes in randomly selected distinct MDR (n=10) and all non-MDR strains (n=10) were studied with respect to housekeeping gene, *recA* (27). Total RNA was extracted from 1 mL of mid-logarithmic bacterial cultures (OD_600_= 0.6) using Nucleospin RNA isolation kit (Macherey-Nagel, Germany). The contaminating DNA was removed by RNase-free DNase I (New England Biolabs, United States). The concentrations and purity of RNA were quantified. Reverse transcription was performed with 2 μg of RNA in accordance to the manufacturer’s instructions using the high-capacity cDNA reverse transcription kit (Applied Biosystems, United Kingdom). Quantification of the expression of the target genes was performed with Power SYBR Green PCR Master Mix (Applied Biosystems, United Kingdom) using the Real-Time T100 Thermal Cycler and Software (Bio-Rad, United States), in triplicate experiments. The C_T_ values of the genes within the study strains were compared. Primers for *cas1* and *cas3* were designed in this study (Table S1) while primer for *recA* was used from a previous study (27).

### 2.6. Whole genome sequencing (WGS) and characterization of complete type I CRISPR-Cas-positive study strains

Total bacterial genome of the representative MDR and non-MDR strains were isolated using Promega Wizard^®^ Genomic DNA Purification Kit. The paired-end libraries were prepared using Nextera XT DNA Library Prep Kit (Illumina, United States) according to manufacturer’s instructions. The genome sequencing was performed using Illumina NovaSeq 6000 with 2x250bp chemistry. The data generated were processed to obtain high-quality reads using TrimGalore (v0.6.7) (quality cutoff Q˃33) and quality checked using FastQC (**v**0.11.3). Reads were assembled using Unicycler (v0.5.0) and annotation was performed using Prokka (v1.14.5), *K. pneumoniae* subsp. *pneumoniae* HS11286 (ASM24018v2) was used as the reference genome for annotation.

### 2.7. Genomic characterization of CRISPR-Cas system possessing genomes

CRISPR-Cas systems in study strains were detected through CRISPRone and spacer sequences were detected through CRISPRTarget website (28) (default cut off score: 20). The default target databases for searching plasmid and phage protospacer sequences in CRISPRTarget were selected as RefSeq and GenBank. Every spacer targeted protospacers (present in invading DNA) with alignment scores ranging from 16-40. Spacers which targeted the protospacer sequences with maximum alignment scores for that particular spacer, was considered for further downstream analysis. Spacers targeting multiple sequences (with maximum alignment scores) were analyzed as per the following protocol-i) targets ranging from 1 to 50: all sequences were analyzed, ii) 51-100 targets: 10% of collection, iii) >100 targets: 5% of collection. For all cases, the first 5 targets analyzed were specifically of *K. pneumoniae* origin, and the rest were randomly selected from the remaining results. Targets of the spacers were backtracked through Entrez nucleotide database to understand the nature of the targeted region i.e., genes present on plasmids, or phage-specific protein, etc. Plasmids targeted by the spacers were downloaded and analysed for the presence of ARGs and replicons in them using ResFinder (v4.6.0) and PlasmidFinder (v2.0.1) respectively. Along with this, the source i.e., organism from where the plasmid was reported, was also noted. Protospacer adjacent motifs (PAMs) i.e. short nucleotide sequences flanking the 5ʹ- and 3ʹ regions of the protospacers were identified using CRISPRTarget.

To correlate CRISPR-Cas system of study strains with global *K. pneumoniae*, a comparative analysis was carried out. For this, CRISPR-CAS++ (https://crisprcas.i2bc.paris-saclay.fr/) was used to collect data on worldwide occurrence of CRISPR-containing *K. pneumoniae* strains with output set as default. Whole genome sequences of the CRISPR-containing global *K. pneumoniae* strains were collected from National Center for Biotechnology Information (NCBI)-Genbank between January 2005 to March 2022 along with their country and source of origin. Genomes (both study and global) were characterized for resistance phenotypes and genotypes using ResFinder (v4.6.0) (https://cge.food.dtu.dk/services/ResFinder/) and sequence types (STs) using Multilocus Sequence Typing (MLST) 2.0 (https://cge.food.dtu.dk/services/MLST/). For the study strains which failed to yield exact STs, genomes were submitted to MLST (https://bigsdb.web.pasteur.fr) and new ST numbers were obtained.

### 2.8. Phylogenetic tree construction

To understand the evolutionary trend of *cas* genes, a phylogenetic tree was prepared. From the fasta files of the whole genome sequences, individual *cas* genes were extracted. The evolutionary history of the individual Cas protein was inferred by Neighbour-Joining method (29) using MEGAXI software package (30) with 1000 bootstrap replication (31). Branches corresponding to partitions reproduced in less than 50% bootstrap replicates are collapsed. The evolutionary distances were computed using the Poisson correction method (32).

### 2.9. Statistical analysis

To understand the correlation between complete type I CRISPR-Cas system and MDR phenotypes in local *K. pneumoniae* strains, two-tailed 2x2 Chi-Square test was performed using from GraphPad (https://www.graphpad.com/quickcalcs/). The Chi-Square values obtained were analysed at 5% level of significance.

## 3. Results

Several relevant aspects of the evaluation of the local study strains were compared with the global strains in the results section. However, for study strains, both laboratory data and genome data were evaluated, whereas for the global strains, all analyses were based on available genomes.

### 3.1. Evaluating CRISPR-Cas systems in study and global *K. pneumoniae* strains

For detection of type I CRISPR-Cas system, presence of *cas1* and *cas3* (signature *cas* genes of type I CRISPR-Cas system) were evaluated by PCR within the study strains (n=241). Seventy-six study strains (32%, 76/241) possessed *cas1* and/or *cas3*, indicating presence of type I CRISPR-Cas systems. However, complete type I CRISPR-Cas system (with both *cas* genes) were found in 25% (60/241) of the total strains, whereas the rest either lacked CRISPR-Cas system (68%, 165/241) or possessed incomplete type I CRISPR-Cas system (7%, 16/241) (Figure S1).

Out of 1613 global *K. pneumoniae* strains, 27% (429/1613) strains had CRISPR elements (possessing either *cas* genes or spacer arrays) in their genome. In this collection, 22% (358/1613) possessed complete type I CRISPR-Cas system and 4% (71/1613) possessed incomplete CRISPR-Cas system with either having truncated *cas* loci (n=9), or containing only CRISPR array (n=62). Both study and global strains showed similar prevalence of complete type I CRISPR-Cas system (25% v/s 22%).

### 3.2. Prevailing trends of CRISPR-Cas system in the study unit, its association with MDR and carbapenem resistance

The study period of thirteen years (2008-2020) was divided into two timeframes - 2008 to 2014 (n=110) and 2015 to 2020 (n=131) for assessing the prevalence of complete type I CRISPR-Cas system over time. Occurrence of complete CRISPR-Cas system was similar between these timeframes with no significant differences in the two timeframes (24%, 26/110 in 2008-2014 v/s 26%, 34/131 in 2015-2020) (Figure S2).

Majority of the study strains (80%, 193/241) exhibited MDR phenotype. Over the years, prevalence of MDR have always been greater than 60% i.e., 90%, 89/110 in 2008-2014 and 79%, 104/131 in 2015-2020, indicating the high load of MDR in study strains (Figure S2). These MDR strains were resistant to carbapenem (last resort drug) and emergence of carbapenem resistance in this unit was noted since 2010. Carbapenem resistance was primarily due to the presence of carbapenemases, namely New Delhi metallo-β-lactamases (*bla*_NDM_). Apart from *bla*_NDM_, these strains harbor different ARGs such as *bla*_CTX-M,TEM,SHV,OXA-1_, *bla*_KPC,OXA-48-like_, *qnrB, qnrS, rmtB,* etc.

Association of MDR with CRISPR-Cas system were assessed by Chi-square test. No significant association was observed between the presence of complete type I CRISPR-Cas system with MDR study strains (Table 1). For the global *K. pneumoniae* (n=1613), similar assessment could not be deduced as AMR phenotypes were only analysed for complete type I CRISPR-positive global strains (n=358), while the comparator group was not analysed.

**Table 1:** Association of complete type I CRISPR-Cas systems with multidrug resistance phenotype and other antimicrobial resistance genes.

| AMR Phenotypes | Complete Type I CRISPR-Cas systems |  |  |
| --- | --- | --- | --- |
|  | Present (n=60) | Absent (n=181) | p-value* |
| **MDR** (n=193) | 50 | 143 | 0.589 |
| **Non-MDR** (n=41) | 10 | 38 |  |
| *bla*<sub>CTX-M</sub> **producers** (n=174) | 45 | 129 | 0.695 |
| *bla*<sub>CTX-M</sub> **non-producers** (n=67) | 15 | 52 |  |
| **Carbapenemase gene producers** (n=119) | 31 | 88 | 0.795 |
| **Carbapenemase non-producers** (n=122) | 29 | 93 |  |
| **PMQR producers** (n=205) | 54 | 151 | 0.303 |
| **PMQR non-producers** (n=36) | 6 | 30 |  |
| **AG resistance gene producers** (n=131) | 34 | 97 | 0.791 |
| **AG resistance gene non-producers** (n=110) | 26 | 84 |  |

Abbreviations: Antimicrobial resistance (AMR); multidrug resistance (MDR); β-lactamase (*bla*), plasmid-mediated quinolone resistance (PMQR), aminoglycoside (AG).
\*Calculated at 5% level of significance.

### 3.3. Antibiotic susceptibility profiling of CRISPR-Cas-positive strains (study and global)

In the study, 60 strains with complete type I CRISPR-Cas system could be divided into MDR (83%, 50/60) and non-MDR (17%, 10/60) (Table 2). The MDR strains exhibited resistance primarily to four groups of antimicrobials *viz.* cephalosporin (100%, 50/50), fluoroquinolone (96%, 49/50), aminoglycoside (96%, 49/50) and carbapenem (64%, 31/50). Non-MDR strains were generally susceptible but some exhibited resistance to cephalosporin (n=5) and fluoroquinolone (n=5) (Table 2).

**Table 2:**
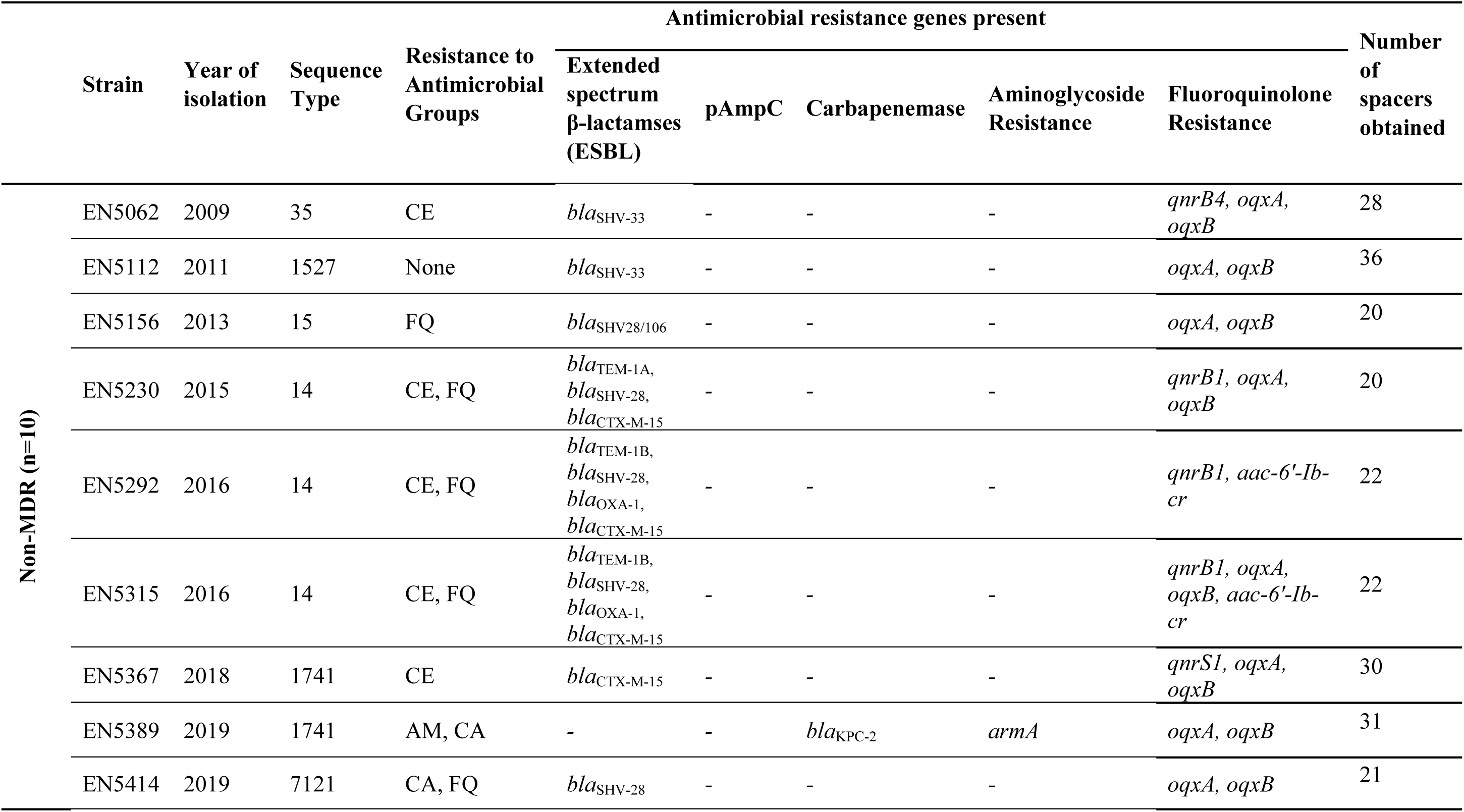
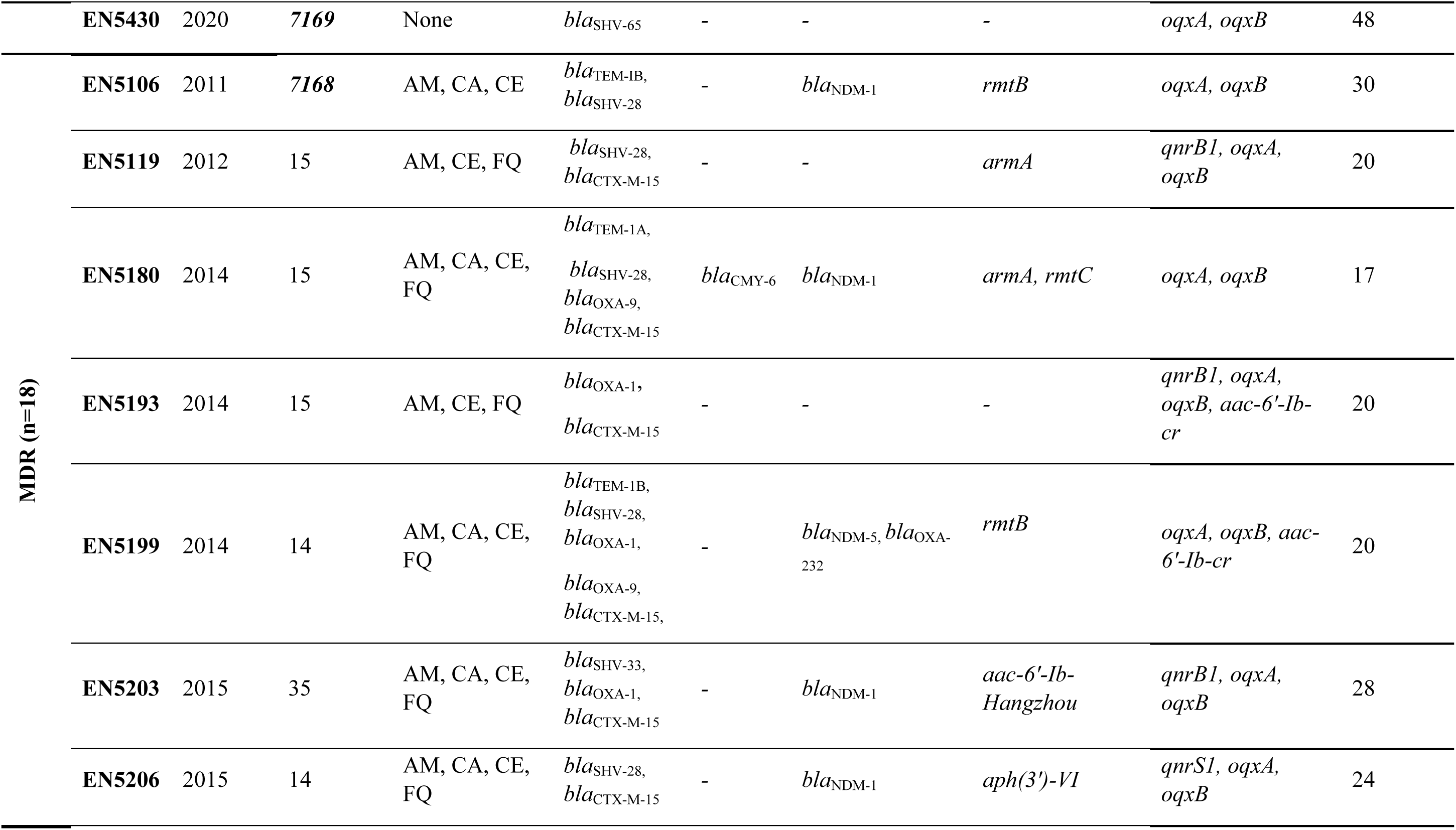
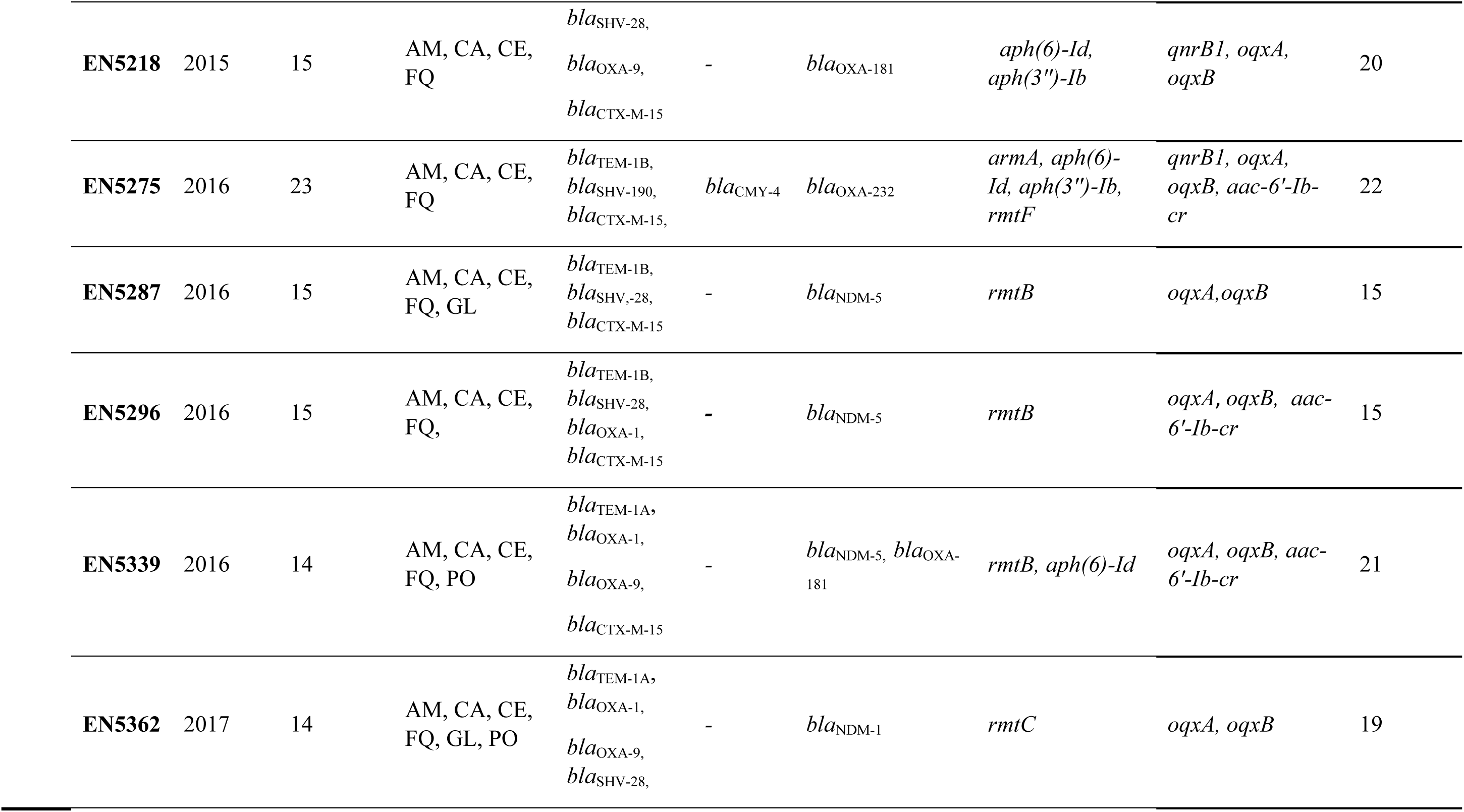
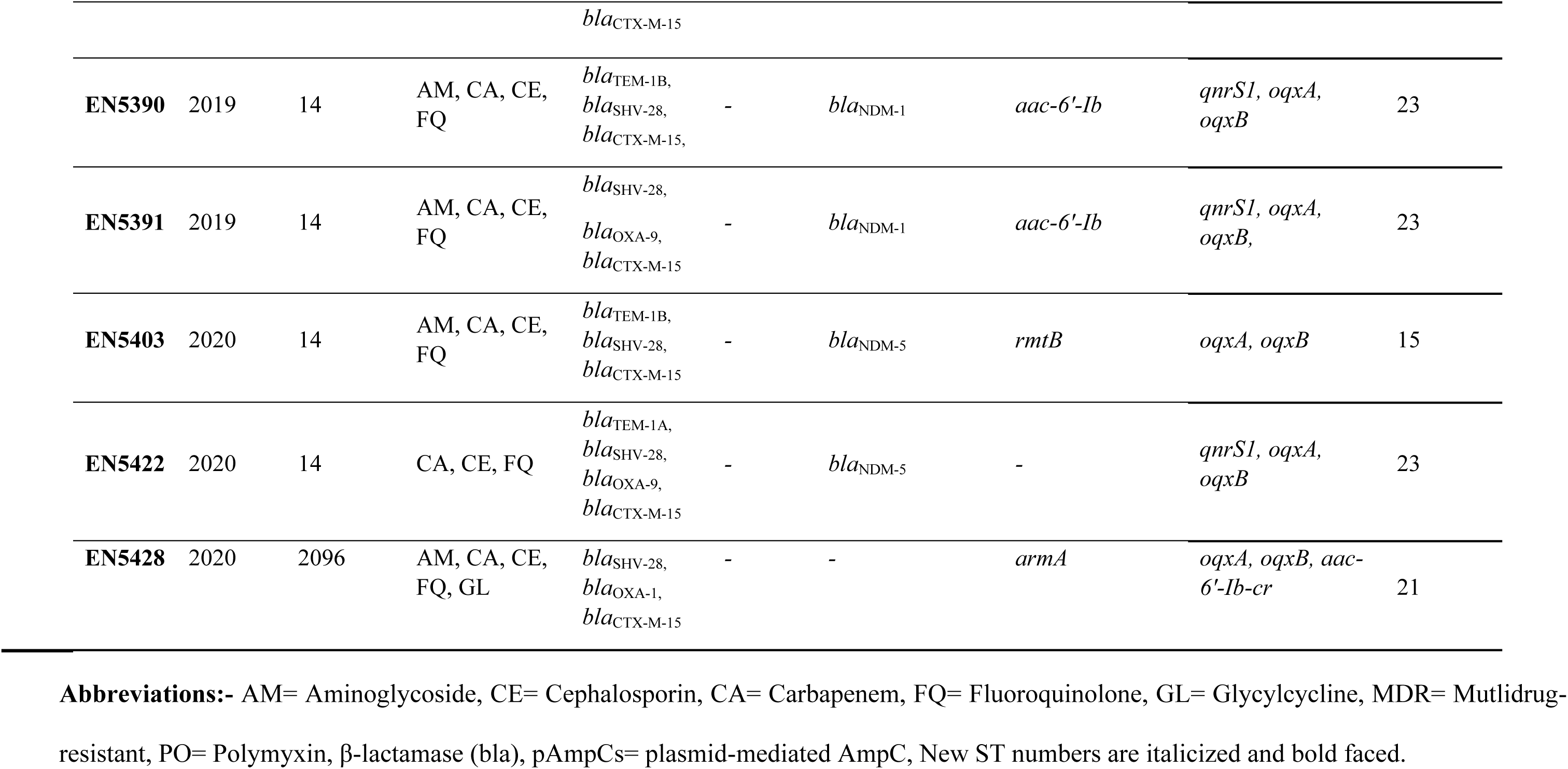
Genomic characterization of type I-E* CRISPR-Cas-possessing *K. pneumoniae* (n=28) from this study.

The global complete type I CRISPR-positive strains were resistant to different classes of β-lactam antimicrobials (80%, 279/358), followed by quinolone (68%, 246/358), aminoglycoside (49%, 176/358) and tetracycline (25%, 90/358) antibiotics. Resistance towards polymyxin (3%, 9/358) group was low. Among type I CRISPR-positive global strains, 47% (168/358) were MDR and 53% (190/358) were non-MDR, based on the phenotypic data obtained from ResFinder (Table S2). Within these non-MDR strains, 20% (73/190) were susceptible to all antimicrobial classes.

### 3.4. Evaluating functionality of CRISPR-Cas system in MDR & non-MDR study strains

The expression of the CRISPR-Cas system was assessed in 20 strains (out of the 60) distributed equally between MDR & non-MDR strains. Out of 50 MDR strains, 10 were randomly selected, however, for non-MDR, all strains (n=10) were included. Mean C_t_ values of housekeeping gene, *recA,* in both group of strains were comparable, 23.77 ± 2.46 in MDR and 23.76 ± 0.96 in non-MDR. Likewise, mean C_t_ values for *cas1* and *cas3* were comparable among MDR and non-MDR, i.e., 28.95 ± 2.15 v/s 28.57 ± 1.17 (*cas1*) and 27.28 ± 2.38 v/s 27.12 ± 1.20 (*cas3*) (Table S3). Though there was no difference between the *cas1* and *cas3* expression in the MDR and the non-MDR group, but, in both groups (MDR and non-MDR) *cas3* was expressed comparatively earlier than *cas1* as indicated by the lower C_t_ values (Table S3).

### 3.5. Resistome analysis of complete type I CRISPR-Cas positive study strains

WGS analysis of all CRISPR-positive non-MDR strains (n=10) was performed; while for the 50 MDR strains, PFGE was carried out as a prerequisite for excluding clonal strains, thereby minimizing selection bias and analyzing only the distinct MDR strains. Thus, 18 CRISPR-positive distinct MDR strains were genome sequenced (Figure S1) and characterized for detailed evaluation of the CRISPR-Cas systems including spacers, PAMs, targeted plasmids, their replicons and resistance determinants. MDR strains (n=18) were primarily resistant towards aminoglycoside, carbapenem, cephalosporin, and fluoroquinolone groups, and possessed diverse ARGs such as ESBLs (*bla*_CTX-M_), pAmpCs (*bla*_CMY_), carbapenemase (*bla*_NDM_), aminoglycoside (*rmtC*) and fluoroquinolone resistance genes (*qnrS* and *qnrB*). On the other hand, non-MDR strains possessed various ESBLs and quinolone resistance genes (*bla*_CTX-M-15_*, bla*_SHV-33,28_, *qnrB1, qnrB4, qnrS1, aac(6ʹ)-Ib-cr*, etc.) except for one non-MDR strain (EN5389), which harbored *bla*_KPC-2_ and *armA* (Table 2). Two non-MDR strains (EN5112 and EN5430) were totally susceptible to all the previously mentioned antimicrobial groups and possessed only *bla*_SHV-33,65_ (Table 2).

### 3.6. Comparative analysis of CRISPR-positive *K. pneumoniae* (study and global) with respect to country of origin, nature and source of isolation

The study strains constitute a localized collection from a single healthcare facility; therefore, the data lack variability with respect to geographic origin, source, and nature of isolation. All study strains were of clinical origin, isolated from the blood of septicemic neonates in India. On the other hand, global strains exhibited diversity in country of origin, nature and source of isolation. Out of the 358 global strains, data on country of origin was available for 316 strains and were reported from 36 countries. Most were concentrated in Asia (n=17) immediately followed by Europe (n=13); while the rest were from North America (n=3), South America (n=2) and Australasia (n=1). More than half of the global strains (55%, 173/316) were reported from three different countries, namely, China (n=84), United States of America (n=50) and India (n=39) (Figure 1, Table S2).

**Figure 1:**
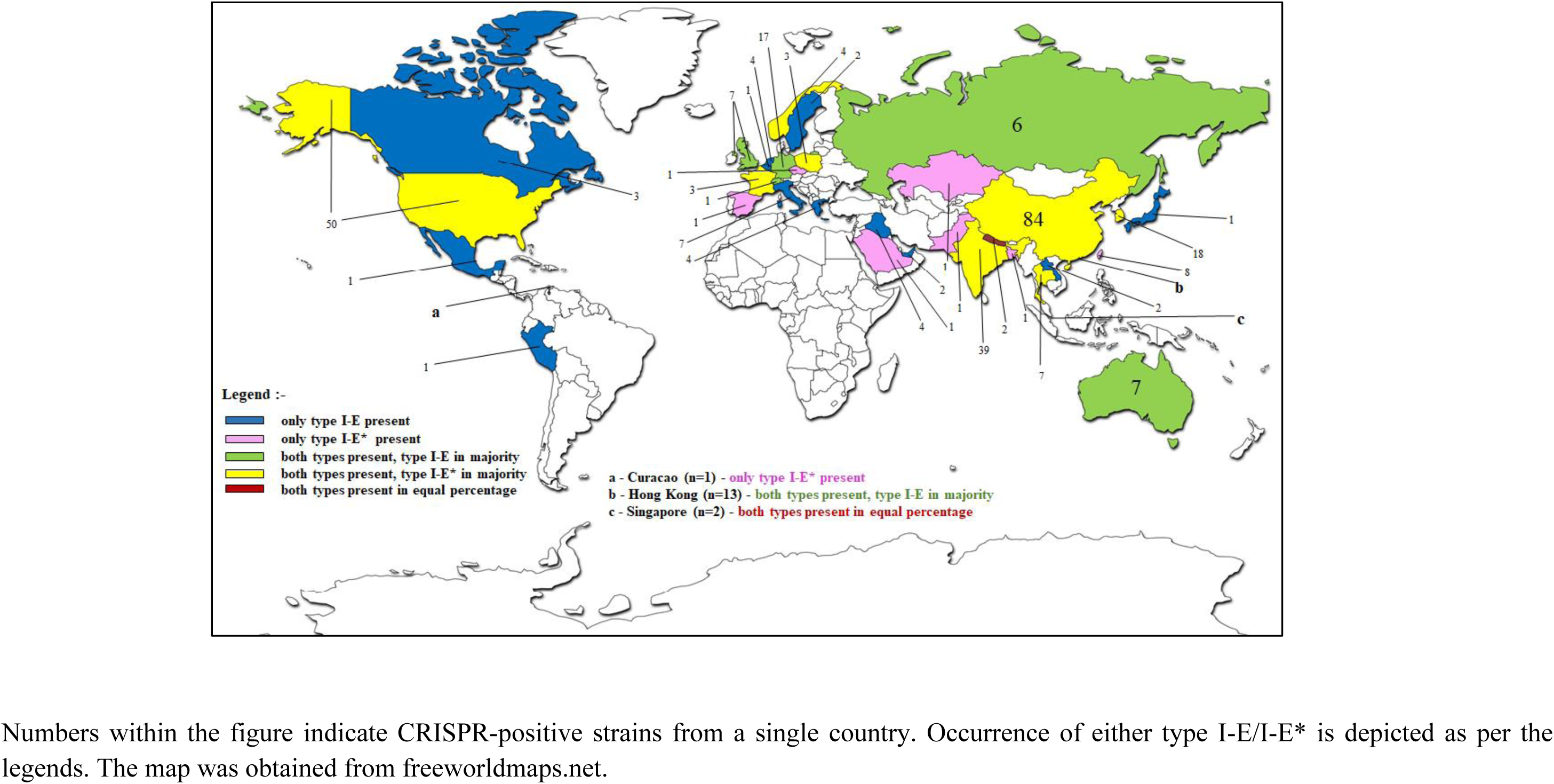
Distribution of type I CRISPR-Cas systems among global *K. pneumoniae* strains (n=316). Numbers within the figure indicate CRISPR-positive strains from a single country. Occurrence of either type I-E/I-E* is depicted as per the legends. The map was obtained from freeworldmaps.net.

Data for source of isolation could be obtained from 314 (88%) global strains, among which clinical strains were predominant (90%, 284/314), followed by those obtained from the environment (n=21). Few strains were isolated from animals (n=7), such as cows, pigs and the Sunda pangolin. All the global clinical strains (n=284) reported were isolated from different organs, tissues and body fluids; most of them (26%, 74/284) being isolated from blood. Strains were also isolated from urine (10%, 28/284) as well as sputum (9%, 26/284) (Figure 1, Table S2).

### 3.7. Characterization and distribution of CRISPR-Cas sub-types among global and local strains

In *K. pneumoniae,* two subtypes of type I CRISPR-Cas systems, type I-E and type I-E*, are known to exist. The study strains (n=28) exhibited only type I-E* systems (Table 2), while the global *K. pneumoniae* (n=1613) showed a nearly equal distribution of type I-E (10%, 154/1613) and I-E* (13%, 204/1613) (Table S2). Distribution of subtypes were different among the continents. Three different trends could be observed- in Asia (n=192) type I-E* was predominant (67%, 129/192) while an opposite trend was noted in Europe (n=61) with type I-E (42/61, 69%) being the prevalent sub-type. A near equal proportion of type I-E* (n=28) and type I-E (n=26) systems were seen in strains from North America (n=54) (Figure 1, Table S2). Among the countries reporting majority of the global CRISPR-positive strains (mentioned in the earlier section), India and China exhibited prevalence of type I-E* CRISPR-Cas systems (79%, 31/39 and 73%, 61/84, respectively) than type I-E systems (21%, 8/39 and 27%, 23/84, respectively). On the contrary, strains from the United States of America (n=50) showed nearly equal distribution of I-E* systems (56%, 28/50) and I-E systems (44%, 22/50) (Figure 1, Table S2).

All the global CRISPR-positive clinical strains were found to possess either type I-E* (58%, 164/284, predominant) or I-E (42%, 120/284) systems. The non-clinical strains (n=29) of environmental and zoological origin, were equally distributed in type I-E (n=13) and type I-E* systems (n=16). Source of origin for few global strains (n=44) could not be recovered, but their presence in both the types were comparable i.e. 20 (I-E) v/s 24 (I-E*). Global strains were primarily reported from blood (n=74) and possessed type I-E* (68%, 50/74) (Table S2), a trend also noted in the local study strains (of blood origin) harboring type I-E*. Non-blood strains on the other hand were equally distributed in both I-E (n=66) and I-E* (n=70) Table S2).

### 3.8. CRISPR-Cas systems and their association with diverse sequence type*s* in study and global *K. pneumoniae*

MLST analysis of the genomic sequences of the study strains (n=28) identified 10 distinct STs, with the non-MDR group exhibiting increased diversity, comprising 7 different STs (Table 2). ST14 (MDR=8/18, non-MDR=3/10), ST15 (MDR=6/18, non-MDR=1/10) and ST35 (MDR=1/18, non-MDR=1/10) were shared between both the MDR and non-MDR groups. Other STs were found either in MDR or non-MDR which included single isolate in ST23, ST2096, ST7168 and ST1527, ST1741, ST7121, ST7169 respectively. ST7168 (n=1) and ST7169 (n=1) were novel STs found in the study strains. ST14 (11/28) and ST15 (7/28) were predominant (Table 2).

For the 358 global strains, STs were successfully determined for 350 strains. For the remaining 8 strains, STs could not be assigned due to gaps in allele sequences or the presence of potentially novel alleles. Global strains were distributed in 65 different STs, belonging to type I-E and I-E* (Table S2). Those possessing type I-E systems predominantly belonged to these STs in the given order-ST147 (n=52) > ST45 (n=14) > ST392 (n=11). On the other hand, strains with type I-E* systems were frequently found in ST23 (n=57) > ST15 (n=45) > ST14 (n=36)> ST35 (n=16), ST2096 (n=8) (Table S2). More than half (51%, 176/350) of the analyzed global strains belonged to the major AMR epidemic clones such as ST14 (n=37), ST15 (n=46), ST35 (n=16), ST45 (n=14), ST147 (n=52) and ST392 (n=11). Some of these epidemic clones (n=68, 19%) such as ST23 (n=57), ST66 (n=3) and ST2096 (n=8) were hypervirulent because of their possession of certain virulence genes different from classical *K pneumoniae*. Type I-E*-bearing strains from India (from global collection) predominantly belonged to ST14 (29%, 9/31), correlating with our local study strain (29%, 8/28). Although a prevalence of type I-E* has been noted from China, but they belonged to ST23 (36%, 29/84) and ST15 (15%, 13/84).

Certain STs (ST14, ST15, ST23 and ST1941) were found in both type I-E and type I-E* global strains. Association of AMR and hypervirulent epidemic clones with CRISPR-Cas positive strains were also noted in the global collection.

### 3.9. Spacer analysis and their variations among study and global strains

The study strains revealed presence of 654 spacers from 28 strains, of which 376 spacers belonged to MDR (18 strains) and 278 to non-MDR (10 strains). Non-MDR strains possessed increased average spacer count compared to MDR strains (28 v/s 21) (Table 3). In the global strains (including both types I-E and I-E* systems), MDR (n=168) showed an increased spacer range and average spacer count (2-86 and 32 respectively) compared to non-MDR strains (n=190) (2–64 and 28.5 respectively). Among the global subtypes, no difference in average spacer count was observed between I-E* MDR (n=83) and non-MDR (n=121) strains (25.1 v/s 25.4). Notably, the average spacer count trends observed in global I-E* strains were consistent with those in the local study strains.

**Table 3:**
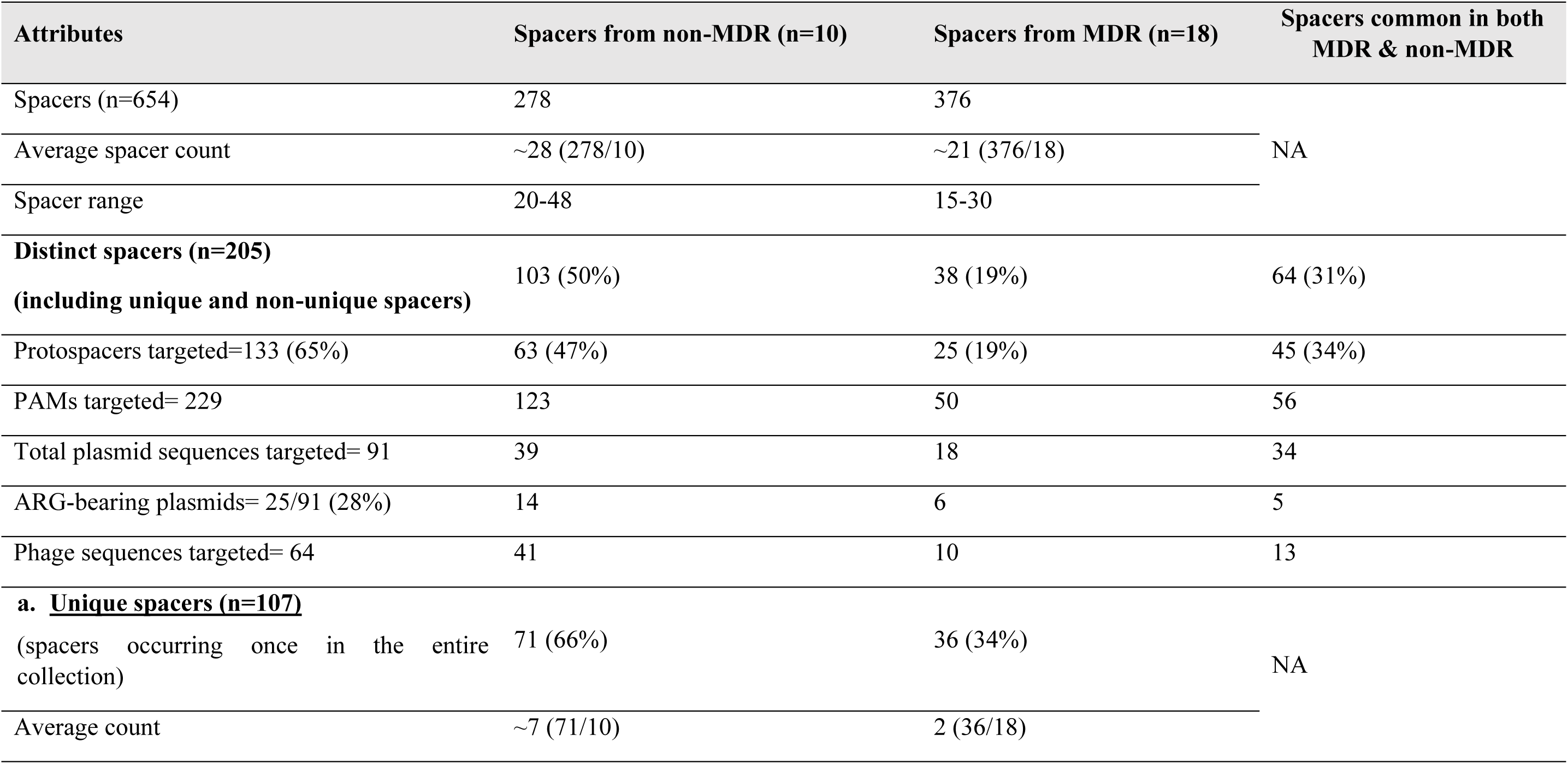
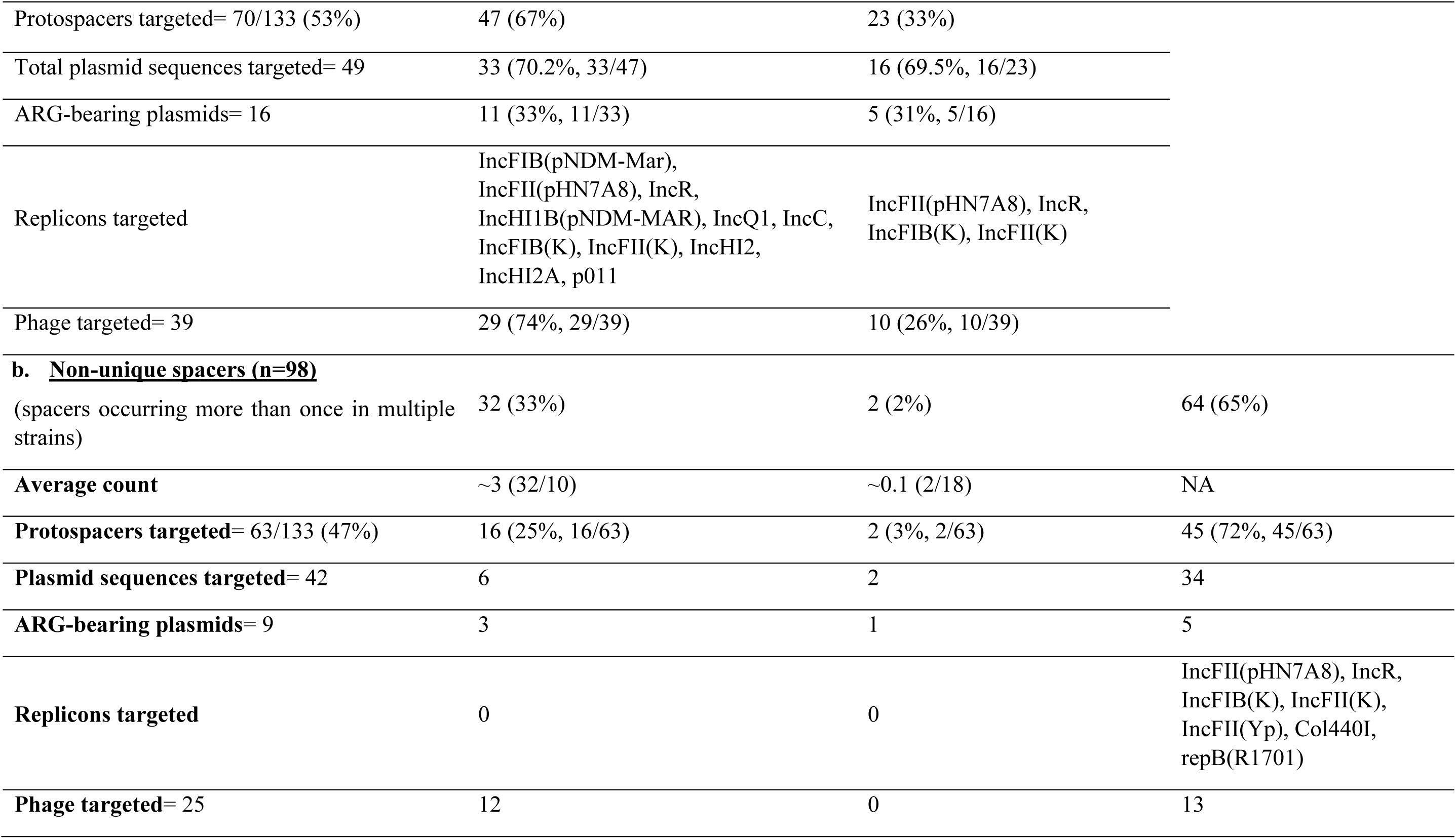
Characterization of spacers in terms of their distribution among the study strains, protospacers targeted, nature of target, carriage of resistance genes and replicons targeted.

| Attributes | Spacers from non-MDR (n=10) | Spacers from MDR (n=18) | Spacers common in both MDR & non-MDR |
| --- | --- | --- | --- |
| Spacers (n=654) | 278 | 376 |  |
| Average spacer count | ~28 (278/10) | ~21 (376/18) | NA |
| Spacer range | 20-48 | 15-30 |  |
| **Distinct spacers (n=205)**<br>**(including unique and non-unique spacers)** | 103 (50%) | 38 (19%) | 64 (31%) |
| Protospacers targeted=133 (65%) | 63 (47%) | 25 (19%) | 45 (34%) |
| PAMs targeted= 229 | 123 | 50 | 56 |
| Total plasmid sequences targeted= 91 | 39 | 18 | 34 |
| ARG-bearing plasmids= 25/91 (28%) | 14 | 6 | 5 |
| Phage sequences targeted= 64 | 41 | 10 | 13 |
| **a. <u>Unique spacers (n=107)</u>**<br>(spacers occurring once in the entire collection) | 71 (66%) | 36 (34%) | NA |
| Average count | ~7 (71/10) | 2 (36/18) |  |
| Protospacers targeted= 70/133 (53%) | 47 (67%) | 23 (33%) |  |
| Total plasmid sequences targeted= 49 | 33 (70.2%, 33/47) | 16 (69.5%, 16/23) |  |
| ARG-bearing plasmids= 16 | 11 (33%, 11/33) | 5 (31%, 5/16) |  |
| Replicons targeted | IncFIB(pNDM-Mar),<br>IncFII(pHN7A8), IncR,<br>IncHI1B(pNDM-MAR), IncQ1, IncC,<br>IncFIB(K), IncFII(K), IncHI2,<br>IncHI2A, p011 | IncFII(pHN7A8), IncR,<br>IncFIB(K), IncFII(K) |  |
| Phage targeted= 39 | 29 (74%, 29/39) | 10 (26%, 10/39) |  |
| **b. <u>Non-unique spacers (n=98)</u>** |  |  |  |
| (spacers occurring more than once in multiple strains) | 32 (33%) | 2 (2%) | 64 (65%) |
| **Average count** | ~3 (32/10) | ~0.1 (2/18) | NA |
| **Protospacers targeted= 63/133 (47%)** | 16 (25%, 16/63) | 2 (3%, 2/63) | 45 (72%, 45/63) |
| **Plasmid sequences targeted= 42** | 6 | 2 | 34 |
| **ARG-bearing plasmids= 9** | 3 | 1 | 5 |
| **Replicons targeted** | 0 | 0 | IncFII(pHN7A8), IncR,<br>IncFIB(K), IncFII(K),<br>IncFII(Yp), Col440I,<br>repB(R1701) |
| **Phage targeted= 25** | 12 | 0 | 13 |

Of the 654 spacers found in the study strains, some spacers appeared once and were designated as unique spacers (n=107), while the remaining (n=574) were found in more than one strain, hence, had duplicates or triplicates across the strains. Of them, 98 spacers (representative of the entire 574 spacers) were denoted as non-unique. These unique (n=107) and non-unique (n=98) spacers altogether formed 205 distinct spacers (Table 3). Individual distribution of spacers corresponding to each strain are depicted in Table S4. These 205 distinct spacers originated from MDR (19%, 38/205), non-MDR (50%, 103/205) and both MDR & non-MDR (31%, 64/205) strains (Table 3, Figures 2.A-B). Targets (protospacers) of these 205 spacers were determined using CRISPRTarget, of which 133 spacers (65%, 133/205) (Table 3) were found to target plasmids and/or phages whereas 72 spacers (35%, 72/205) did not yield any result (Tables 4.A-B, S4). Among these 133 spacers which yielded results, 69 (52%, 69/133) targeted plasmid sequences while 42 (32%, 42/133) targeted phage sequences, and 22 spacers targeted both plasmid and phage sequences (17, 22/133) (Tables 3, S4 and Figures 2.A-B). Plasmid sequences targeted by spacers from non-MDR were high compared to MDRs (39 v/s 18) (Table 3).

**Figure 2:**
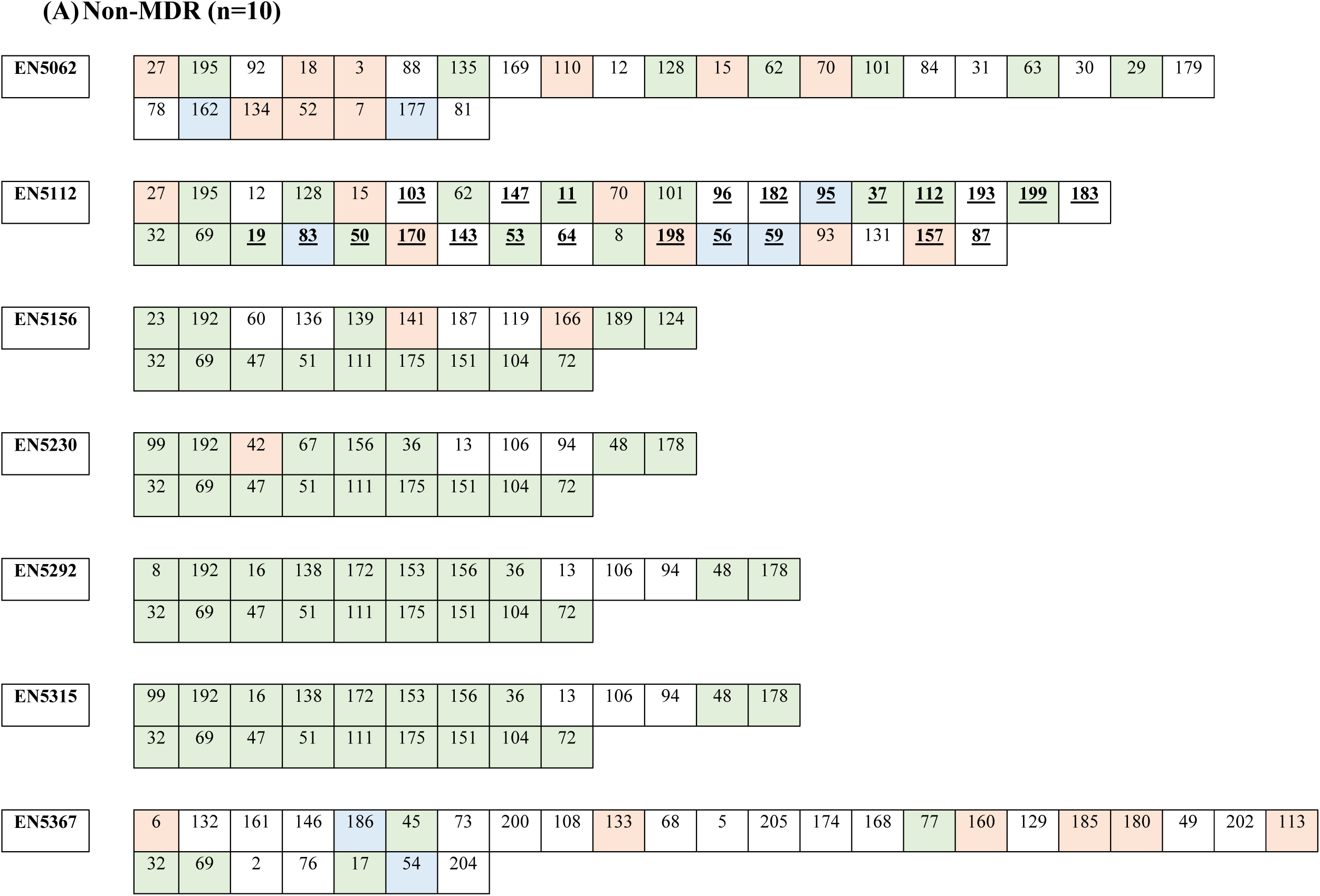
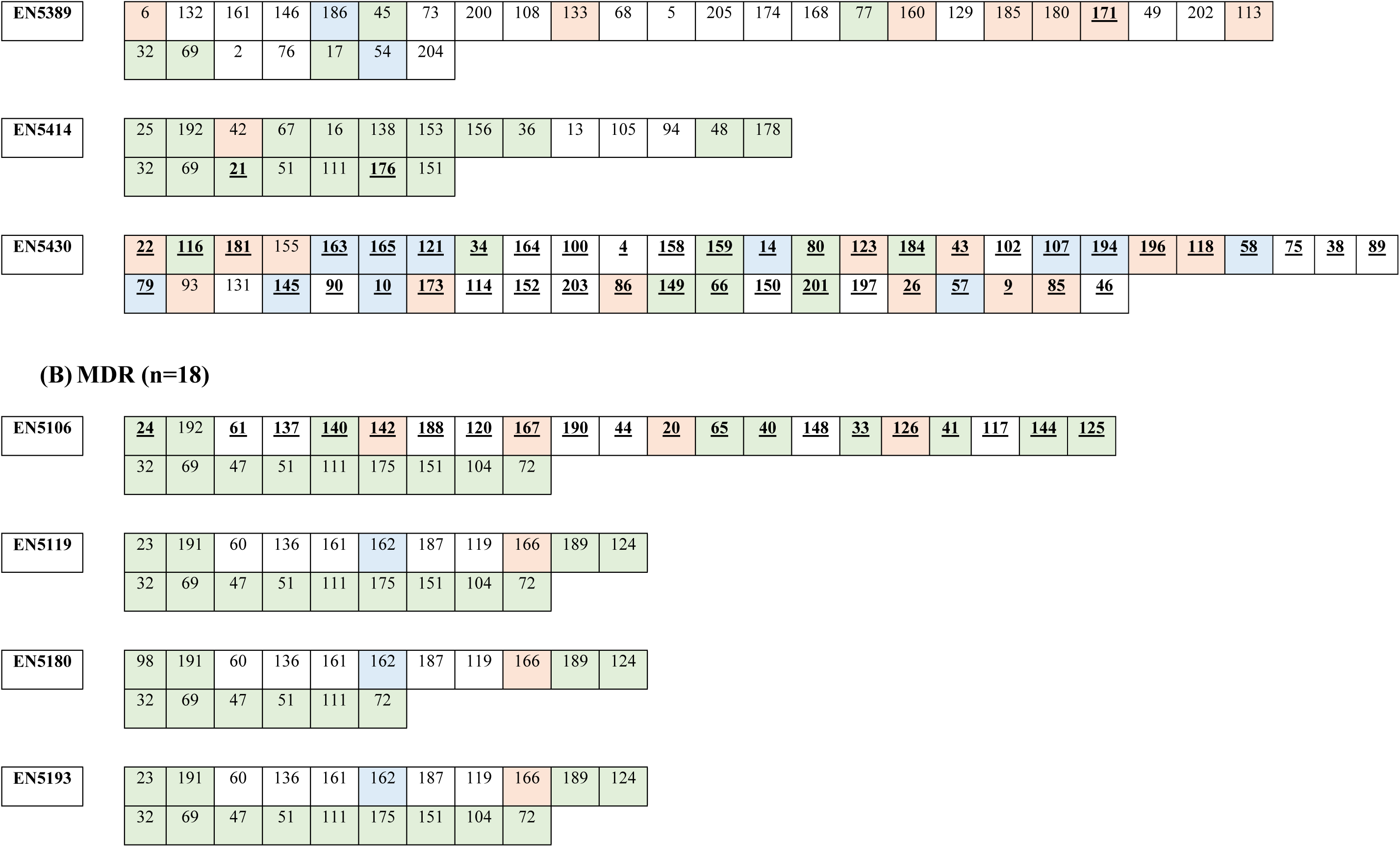
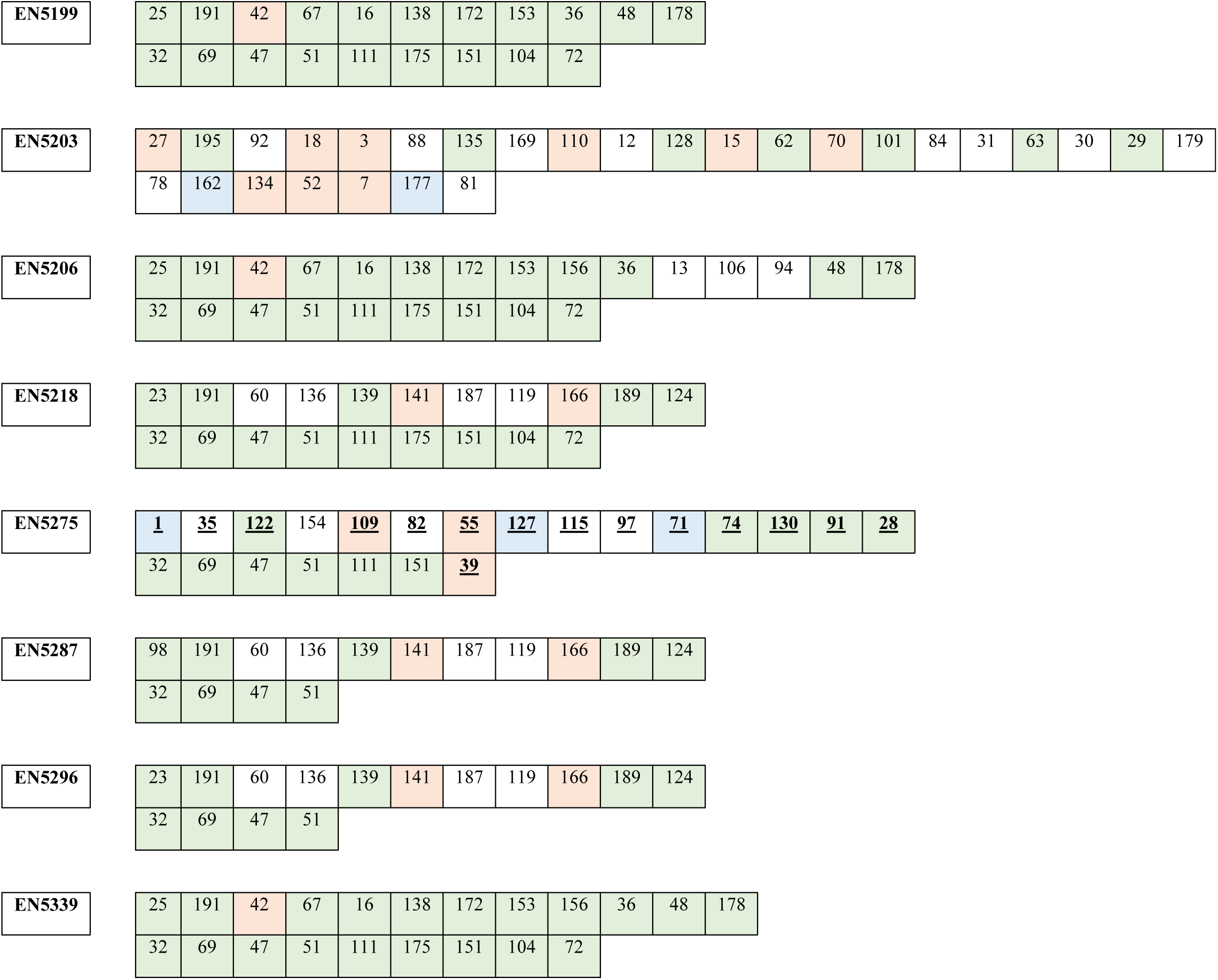
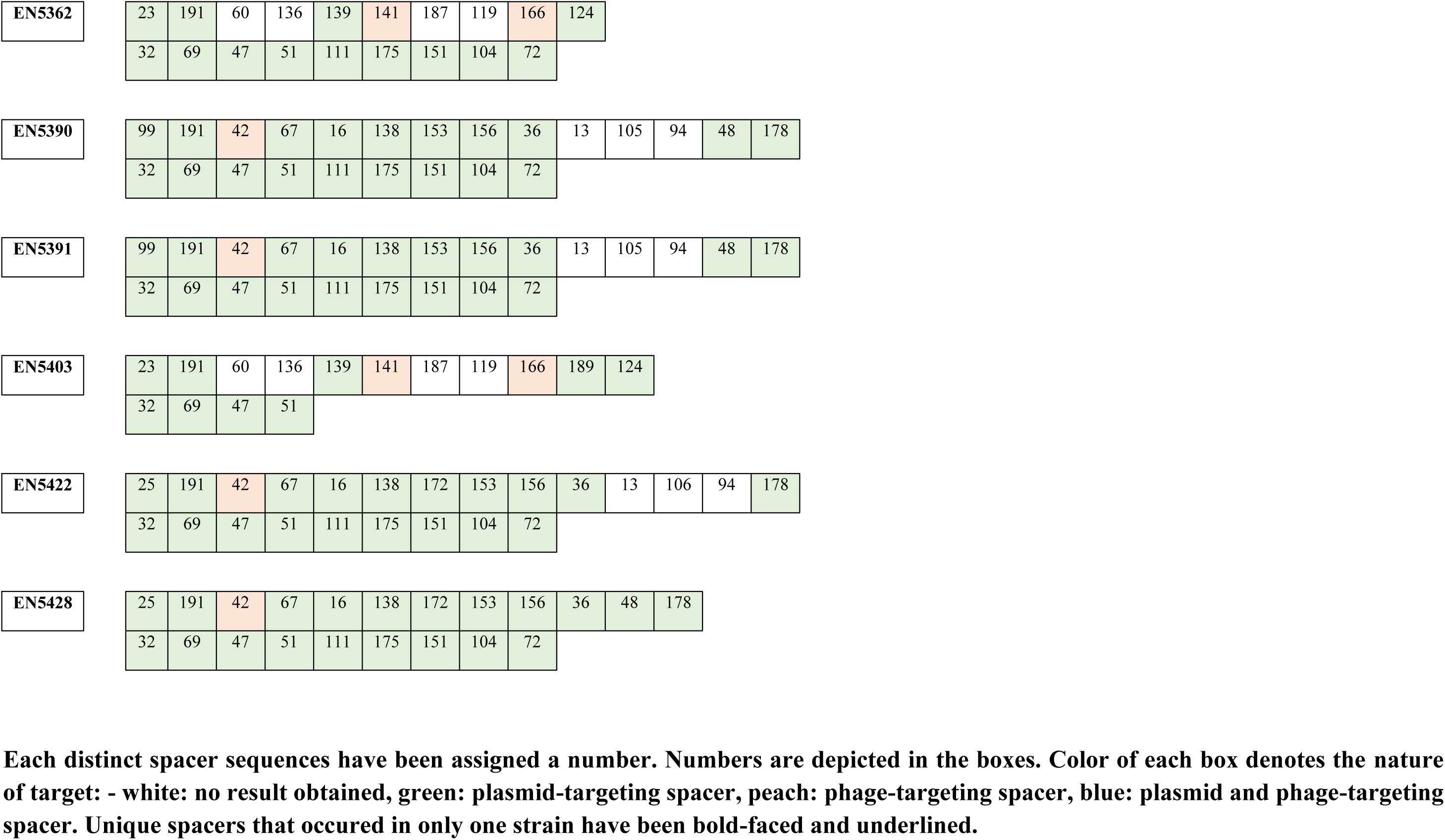
Characterization of distinct spacers (n=205) retrieved from type I-E* CRISPR-Cas-harbouring study strains in terms of their occurrences and nature of target.

Non-MDR strains possessed more unique spacers (66%, 71/107) compared to MDR strains (34%, 36/107). A striking difference in average number of unique spacers was noted in non-MDR: 7.1 (71/10) v/s MDR: 2 (36/18) (Table 3), indicating increased diversity of spacers in non-MDR compared to MDR strains. Protospacers could be identified for 70 unique spacers, which targeted plasmids (n=49) and phages (n=39). Amongst them 18 spacers targeted both plasmid and phage sequences (Tables 3, S4, Figures 2.A-B). Unique spacers derived from non-MDR strains (67%, 49/70) targeted more protospacers compared to those from MDR strains (33%, 23/70) (Tables 3, S4).

CRISPR-positive global strains analyzed in this study revealed presence of 10,795 spacers (Table S2). Spacers from these strains (n=358) targeted plasmids and phages. Among them, 9 strains harbored spacers which targeted plasmids only. Most of these plasmids were from *Klebsiella* sp. These plasmids harbor different ARGs such as *bla*_KPC_, *bla*_TEM-1_, *dfrA14, qnrB1, aph*(*6*)*-Id, aph(3’’)-Ib, sul2*, etc. Plasmids targeted by these spacers belonged to IncFIA(HI1), IncFIB, IncFIB(K), IncFII, IncC replicons (Table S2).

On the other hand, 3 global strains i.e. FDAARGOS_1312, Kp1 and KPHS1249 predominantly consisted of phage target sites in their spacer array, all of which targeted *Klebsiella* phages (Table S2).

### 3.10. Analysis of plasmid-targeting spacers: targeted ARGs and replicons

Plasmid-targeting spacers (n=91) i.e. spacers targeting only plasmids (n=69) as well as those targeting both plasmids and phages (n=22) targeted 263 plasmids, of which 133 plasmids were non-duplicate. Plasmid-targeting spacers were found to be higher in non-MDR strains (n=73) compared to MDR strains (n=52) (Tables 3, S4). These plasmids were found in 43 diverse bacteria belonging to 13 various orders (Figure 3). Majority of the plasmids were retrieved from Enterobacterales (n=12), followed by Burkholderiales (n=7) and Sphingomonadales (n=6). Among Enterobacterales, plasmids from five different species of *Klebsiella (K. grimontii, K. michiganensis, K. pneumoniae, K. quasipneumoniae* and *K. variicola*) along with *Enterobacter cloacae complex, Escherichia coli*, *Pantoea deleyi, Raoultella ornitholytica, Salmonella enterica, Serratia mercescens,* and *Shigella boydii* were targeted (Figure 3, Table S5). Plasmid-targeting spacers primarily targeted plasmid sequences from Enterobacterales, except for one spacer (spacer no. 116) which targeted protospacer of non-Enterobacterales origin (Table 4.A). These spacers mostly targeted direct repeat regions (26%, 24/91) and hypothetical proteins (18%, 16/91) (Tables 4.A-B and S5). Additionally, some of these spacers could target genes encoding metabolic enzymes like DNA cytosine methyltranferase (n=3), phage-specific proteins like phage portal proteins involved in capsid assembly (n=2). Other genes encoding various proteins were also targeted such as those encoding TolC family protein (outer membrane efflux proteins found in Gram-negative bacteria), integrase, and ParB/RepB/Spo0J family partition protein.

**Figure 3:**
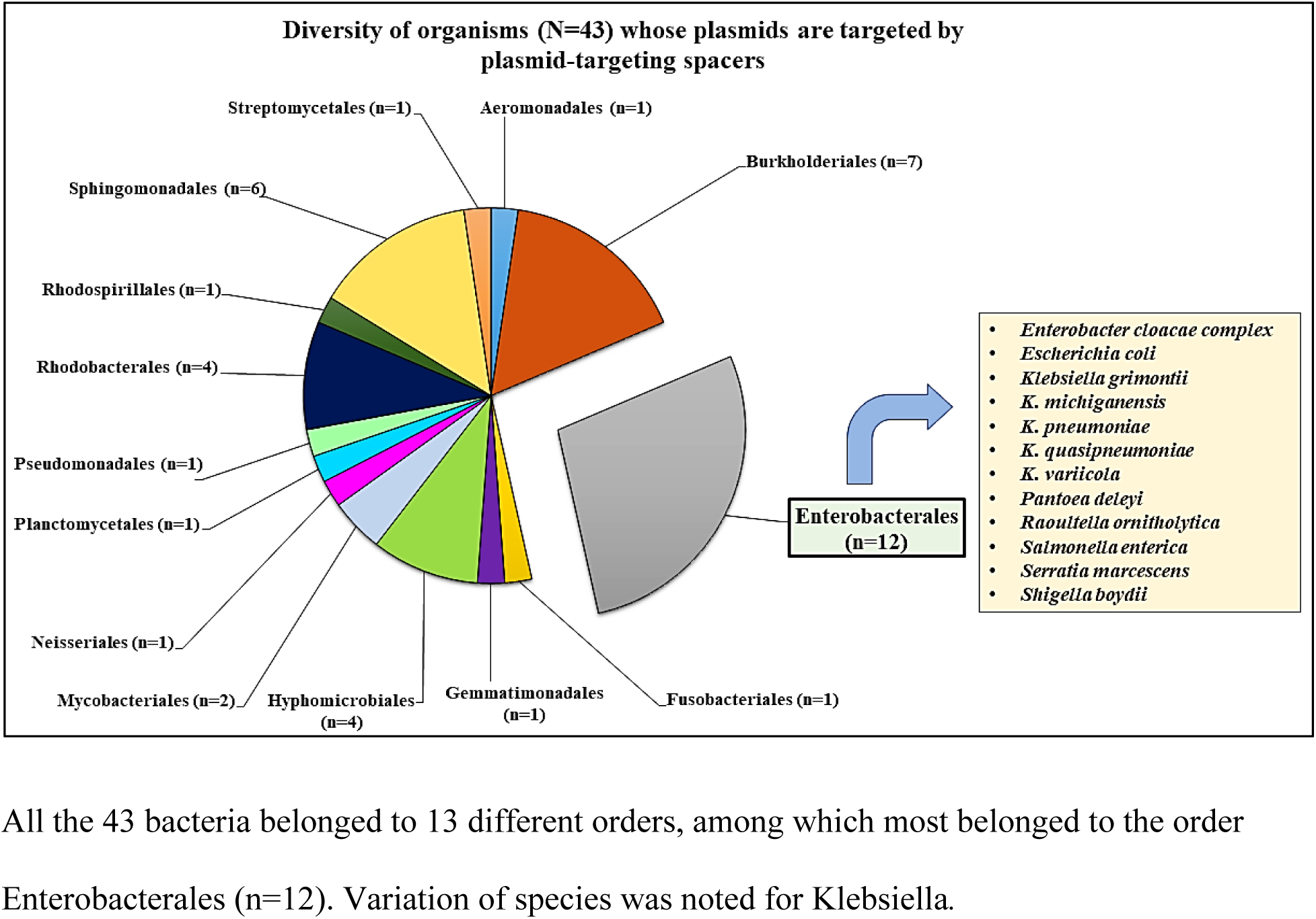
Origin of plasmids targeted by plasmid-targeting spacers of this study. All the 43 bacteria belonged to 13 different orders, among which most belonged to the order Enterobacterales (n=12). Variation of species was noted for Klebsiella.

**Table 4:**
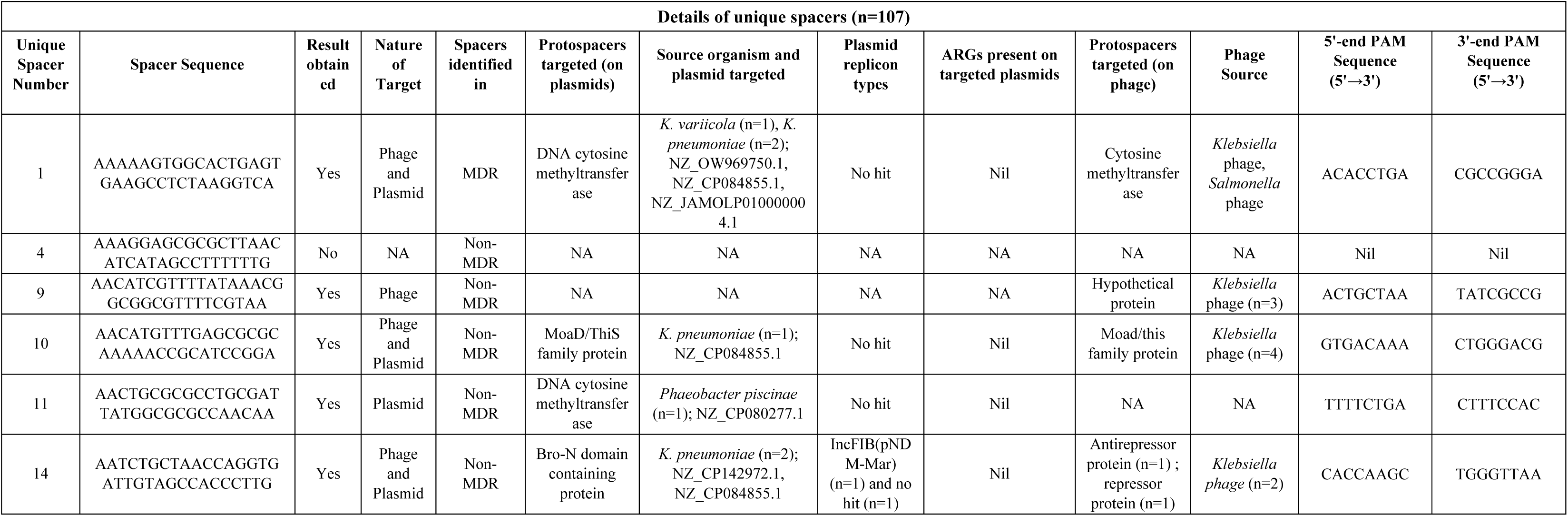
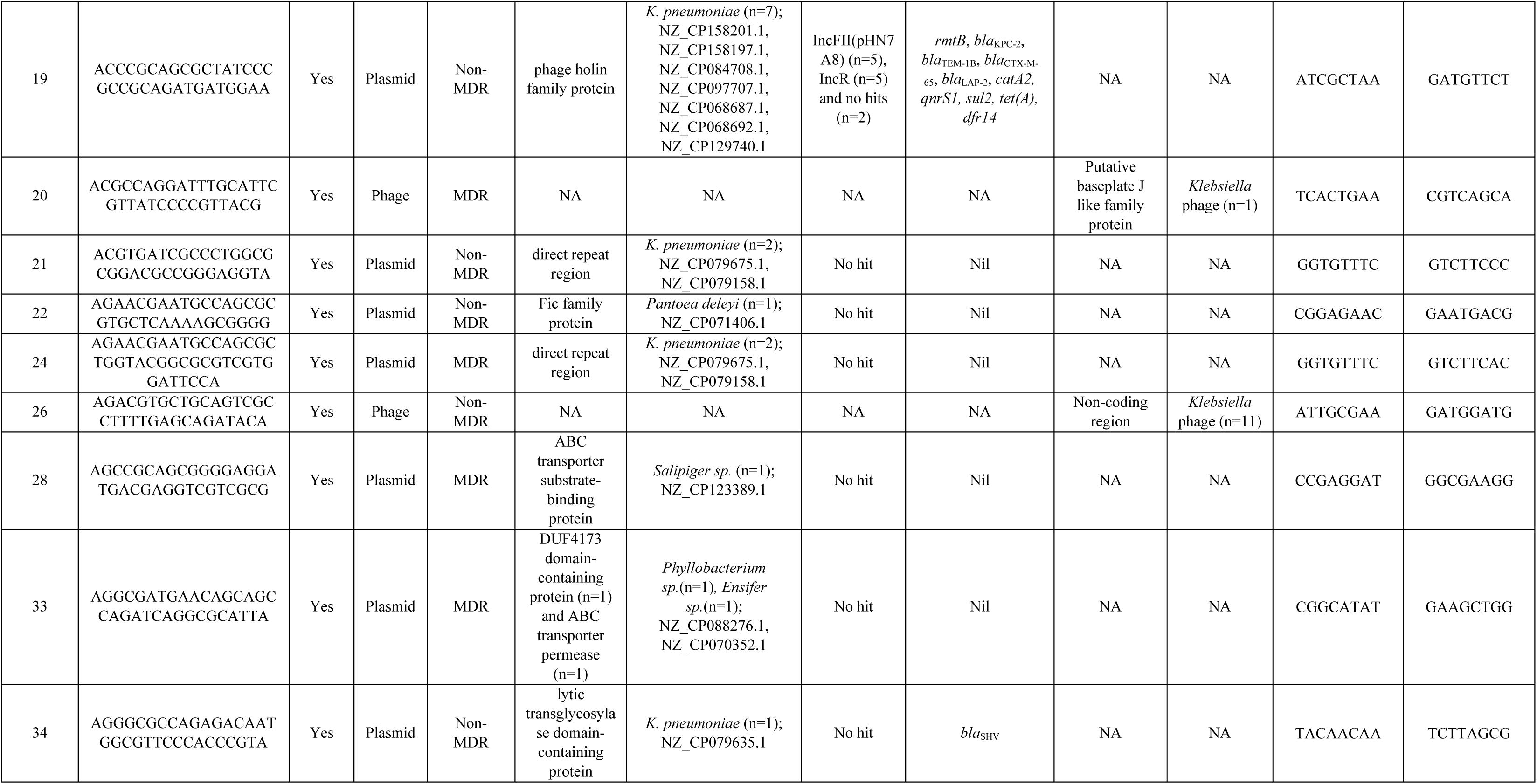
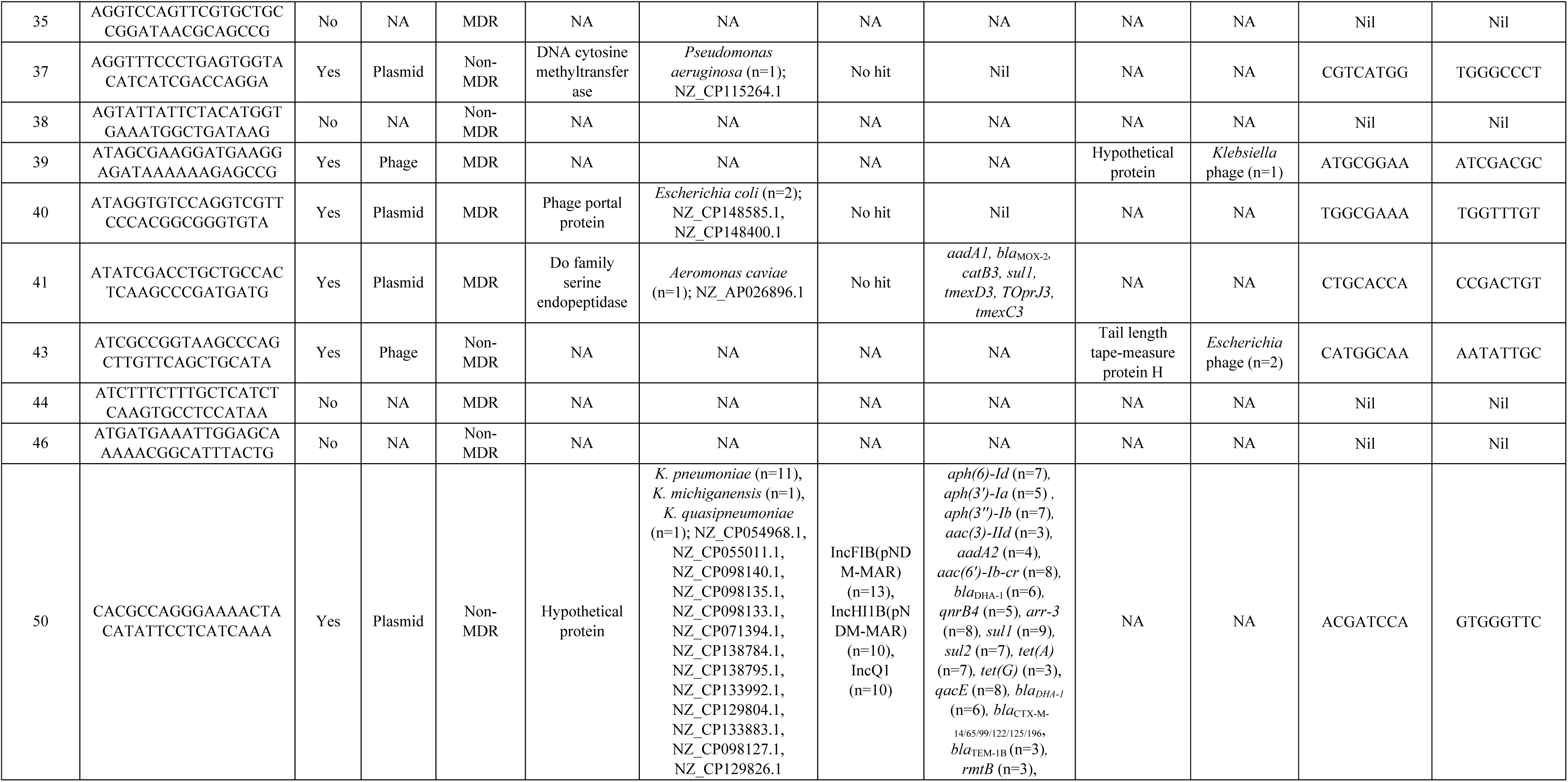
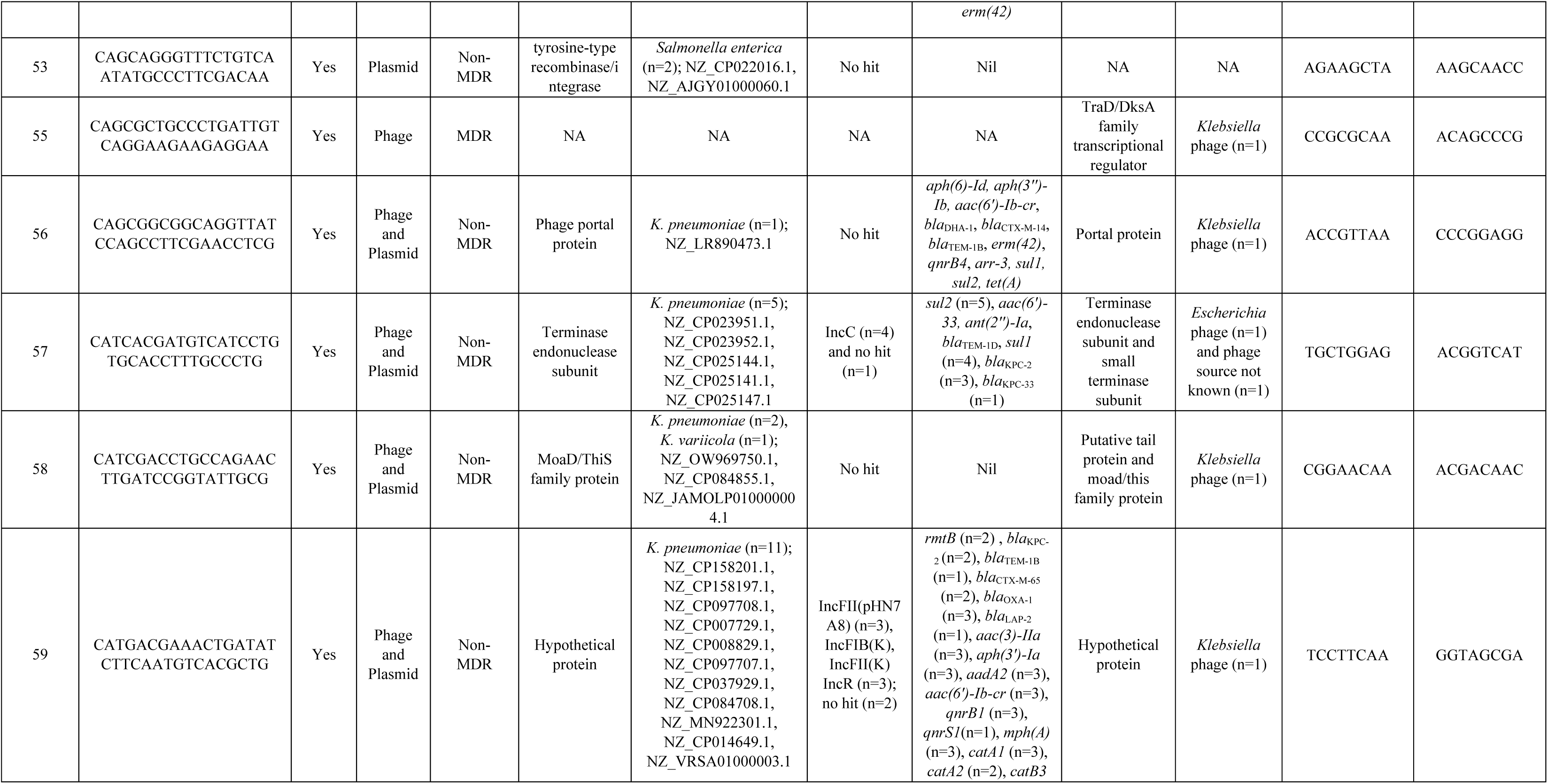
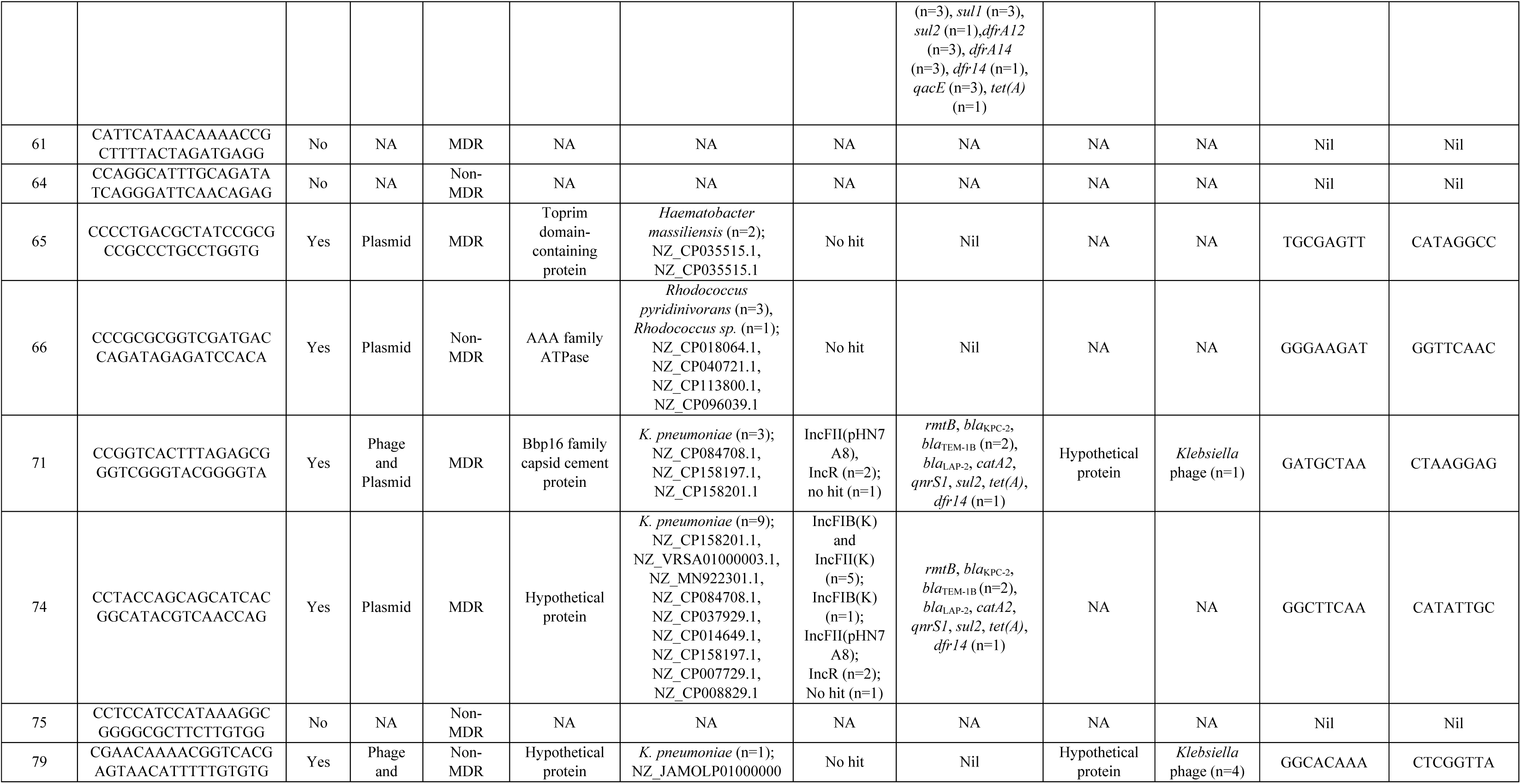
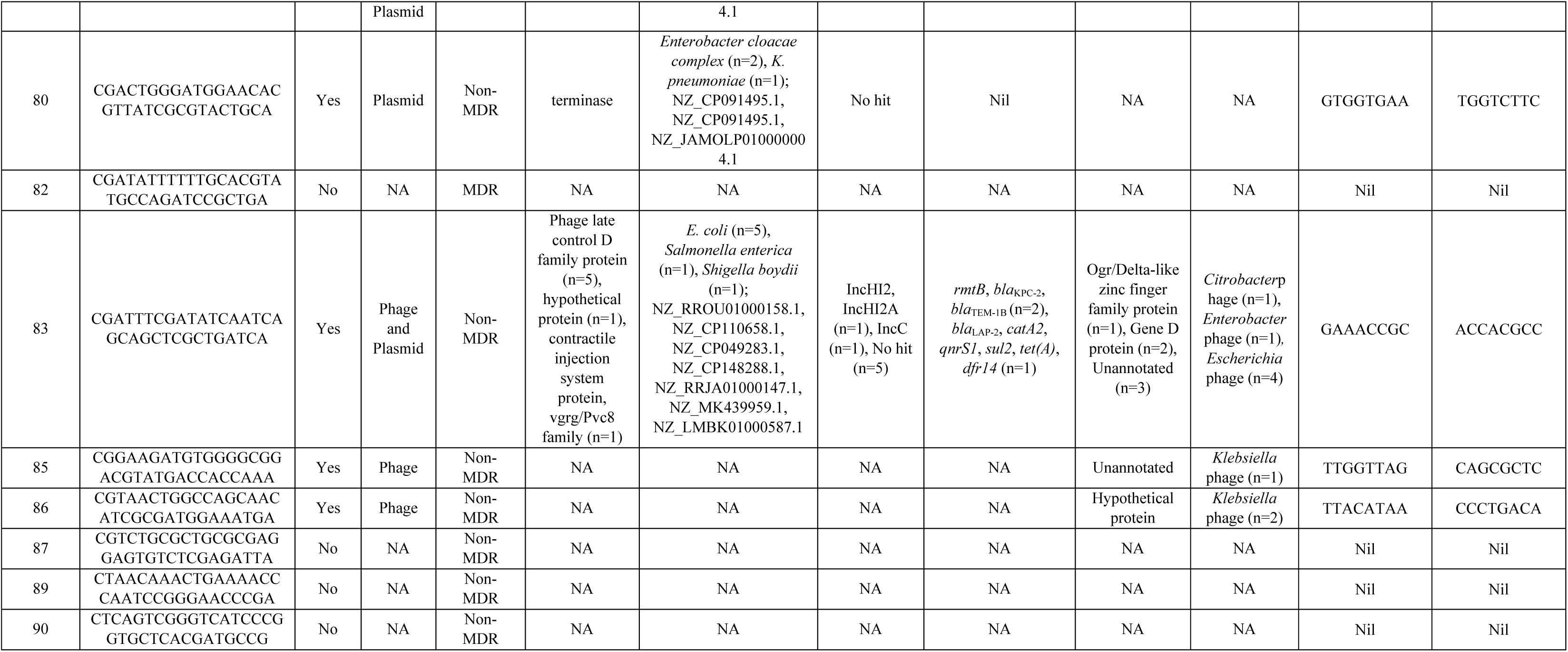
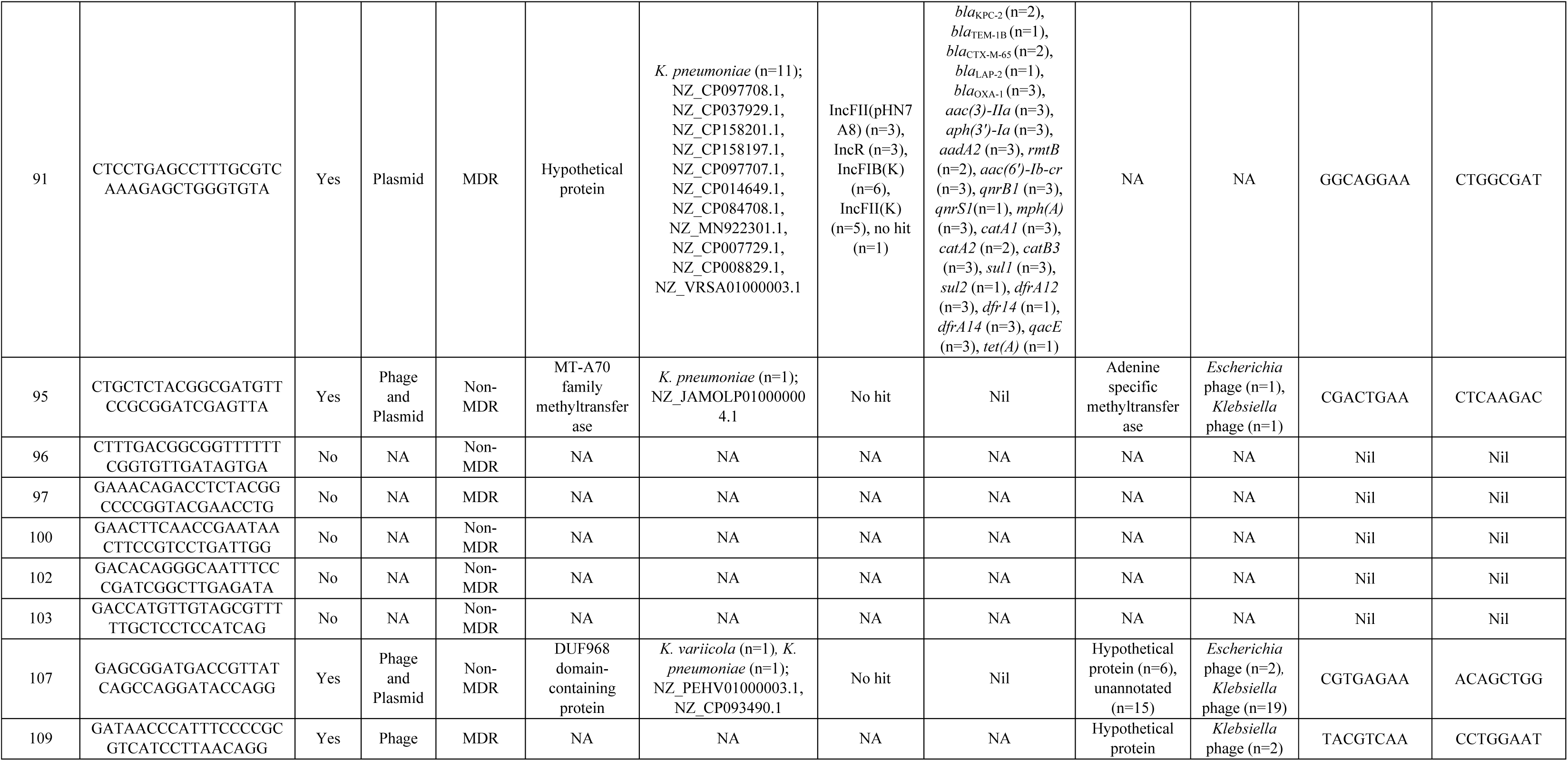
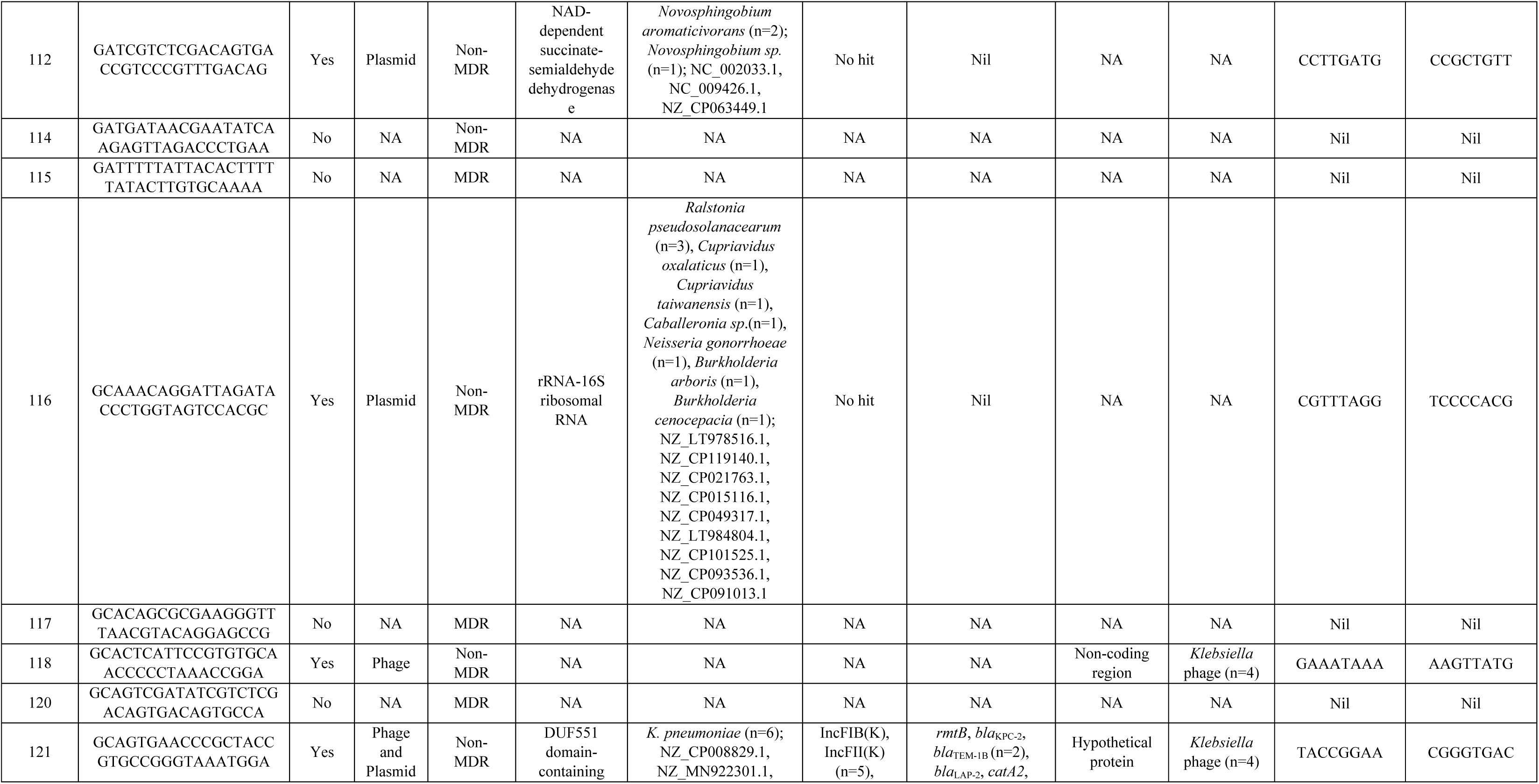
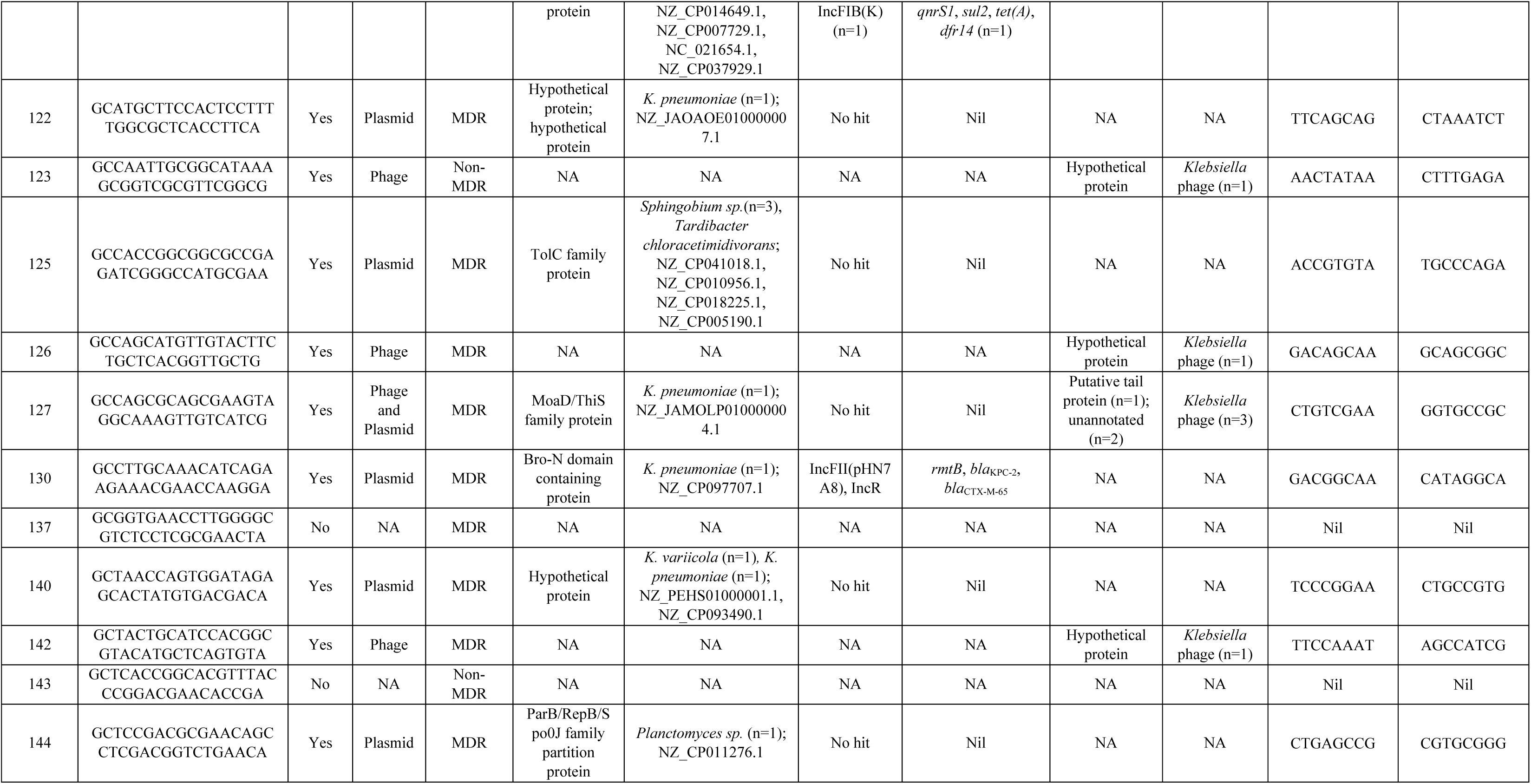
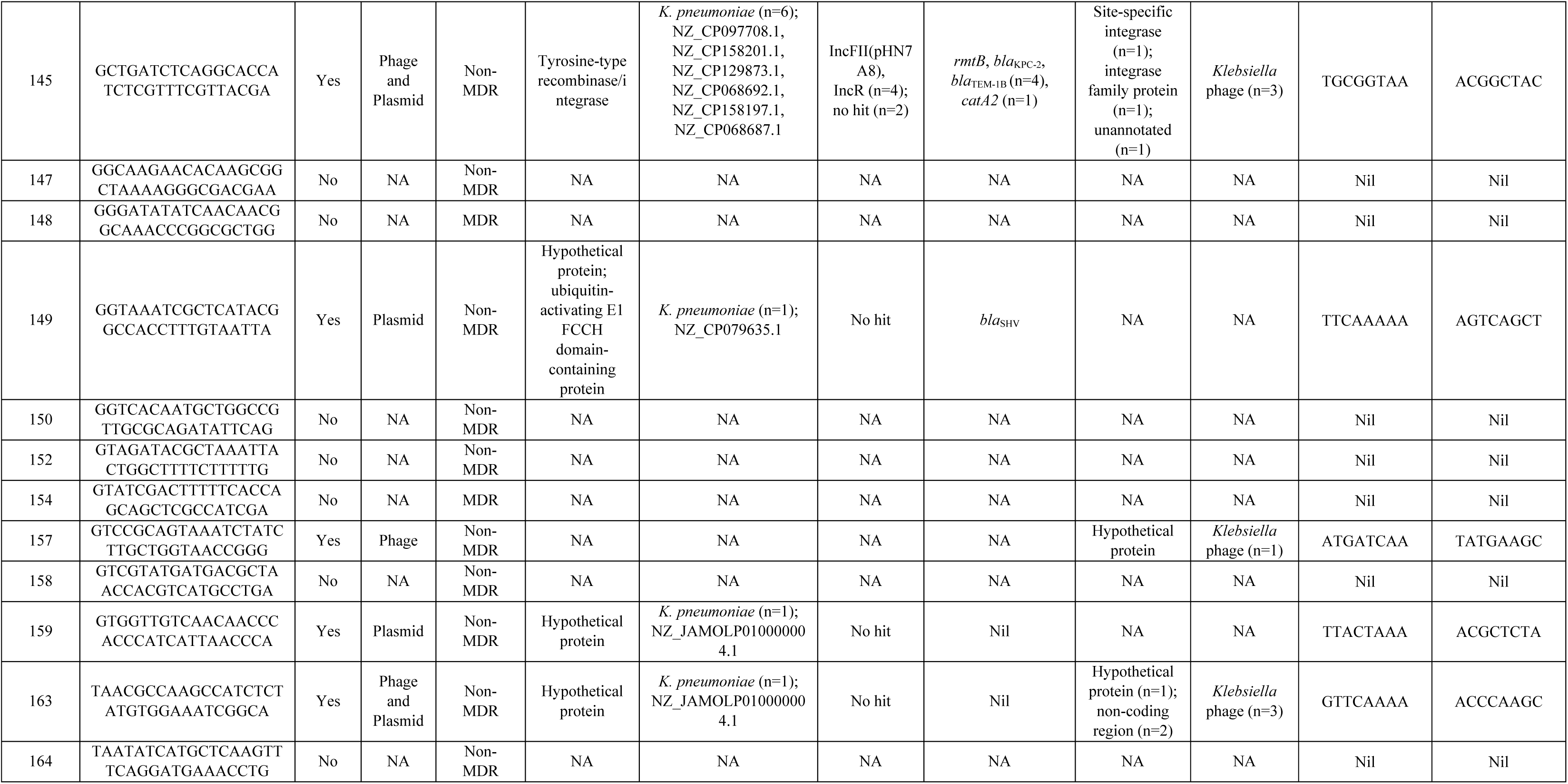
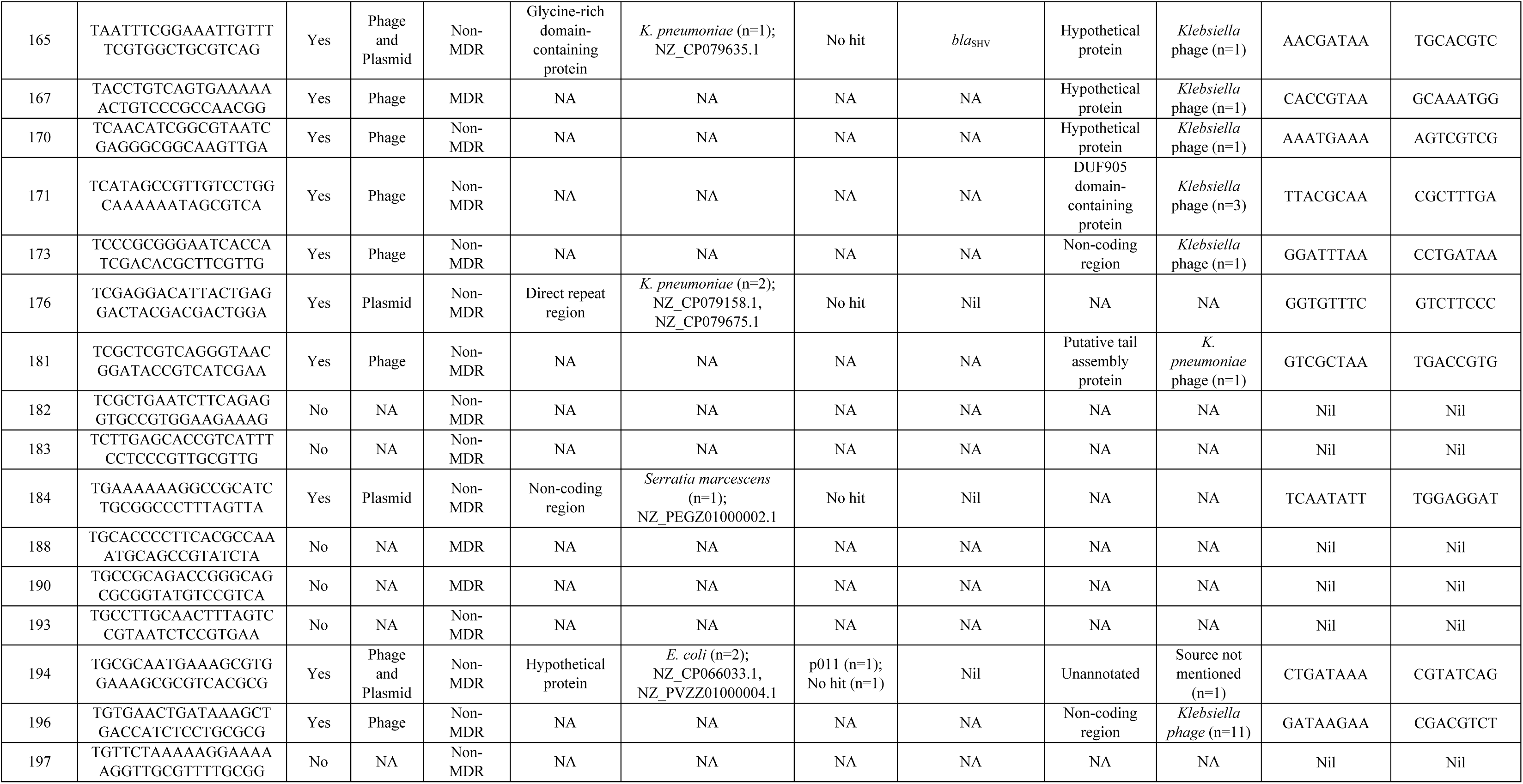
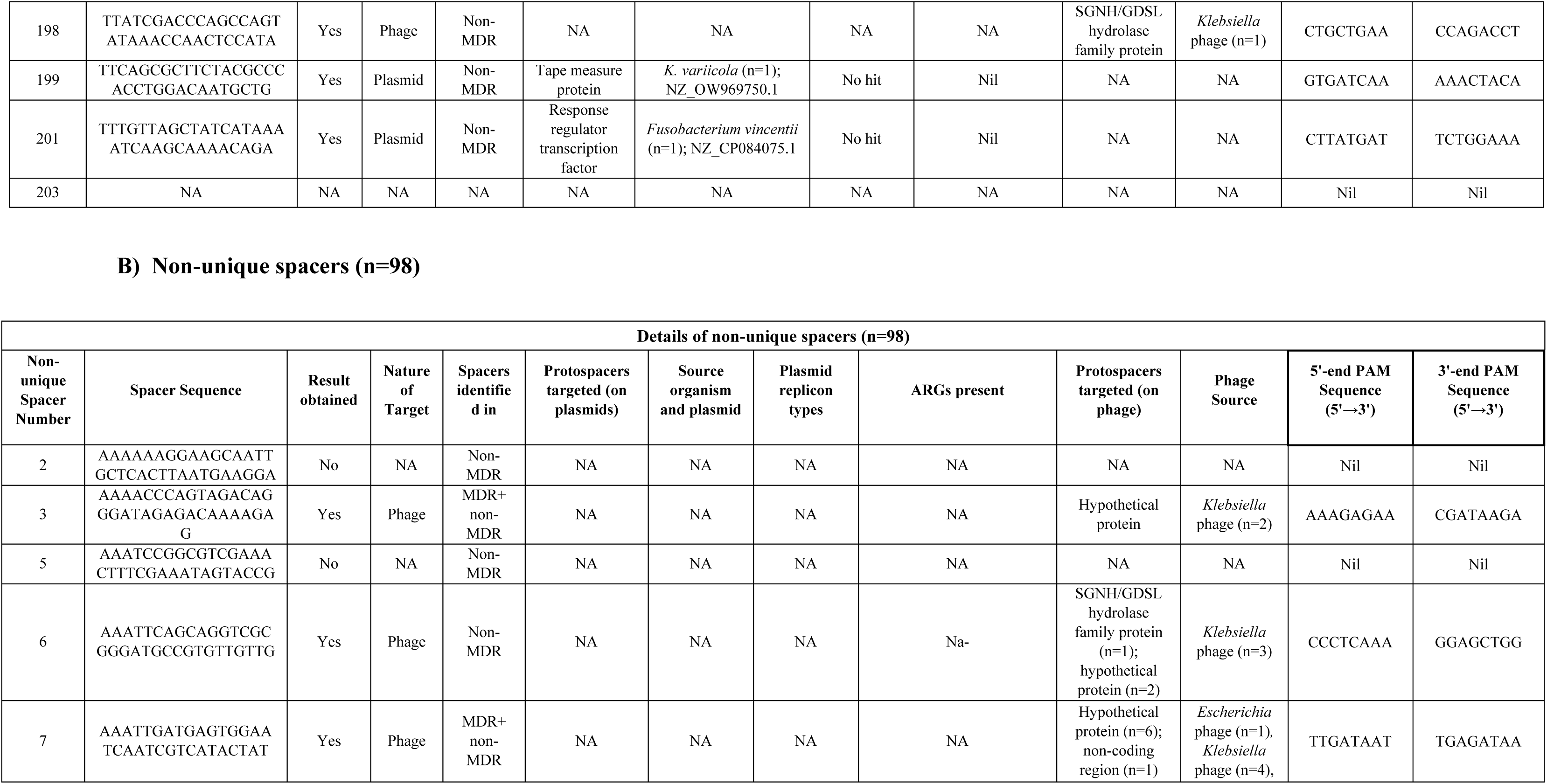
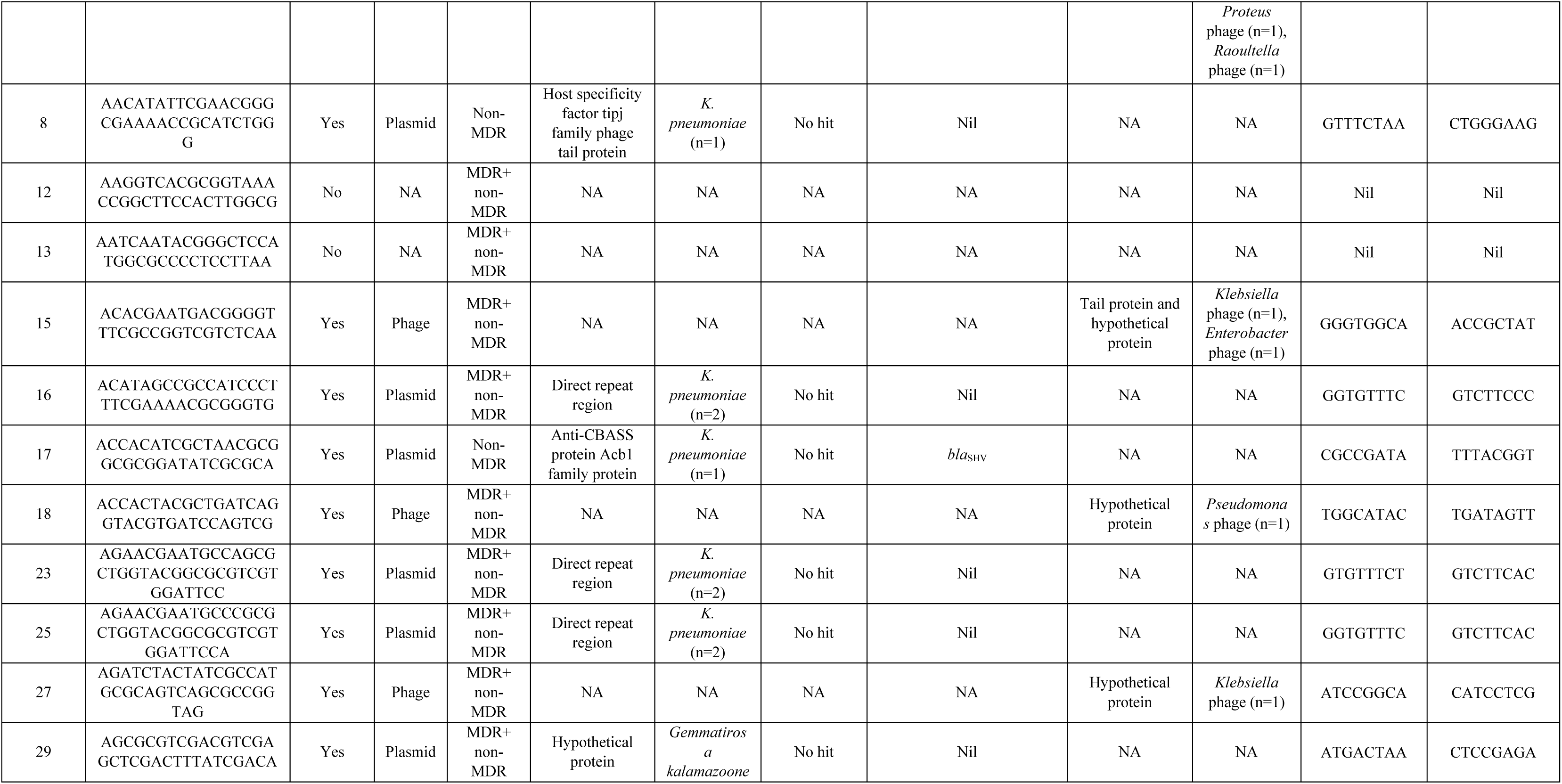
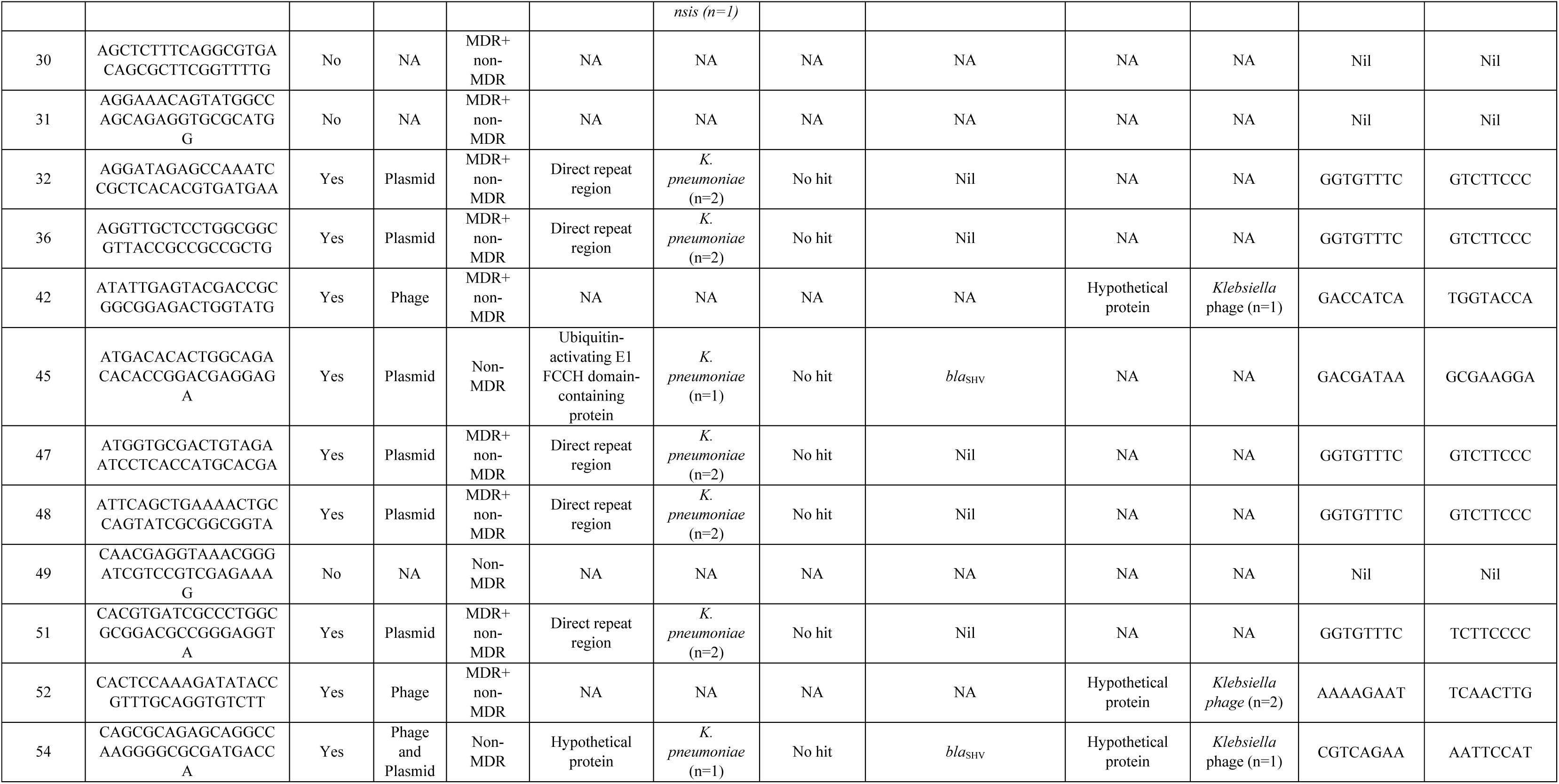
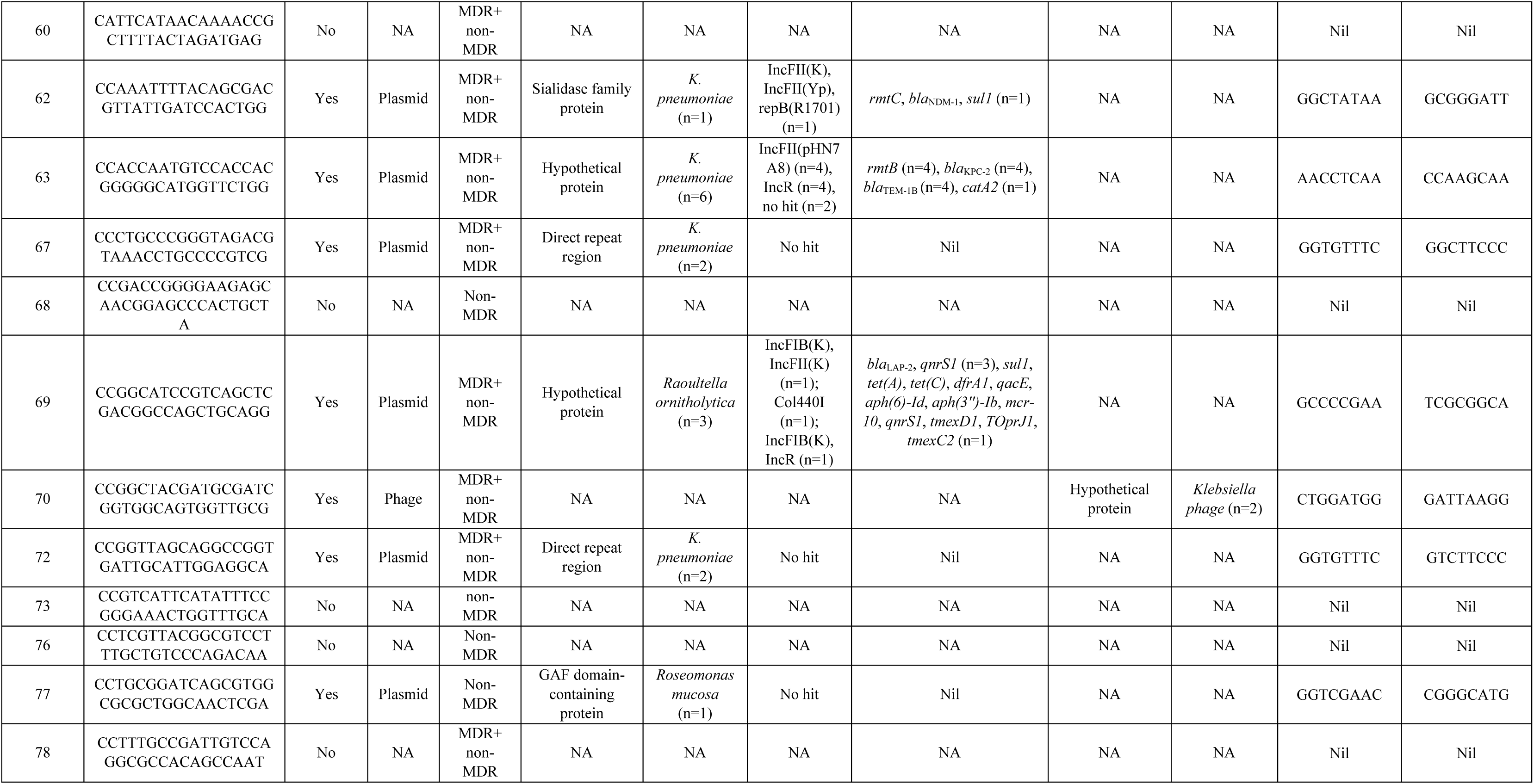
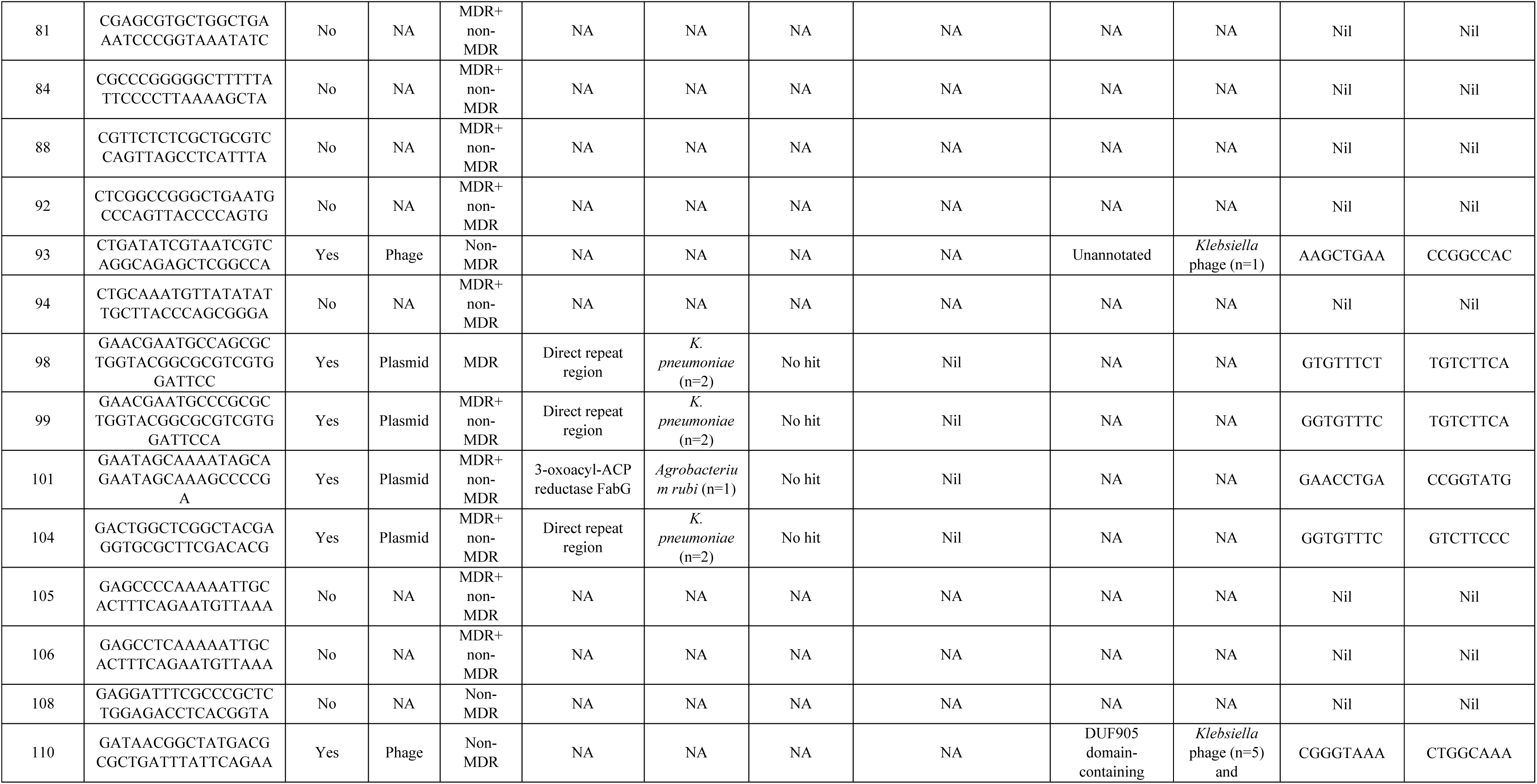
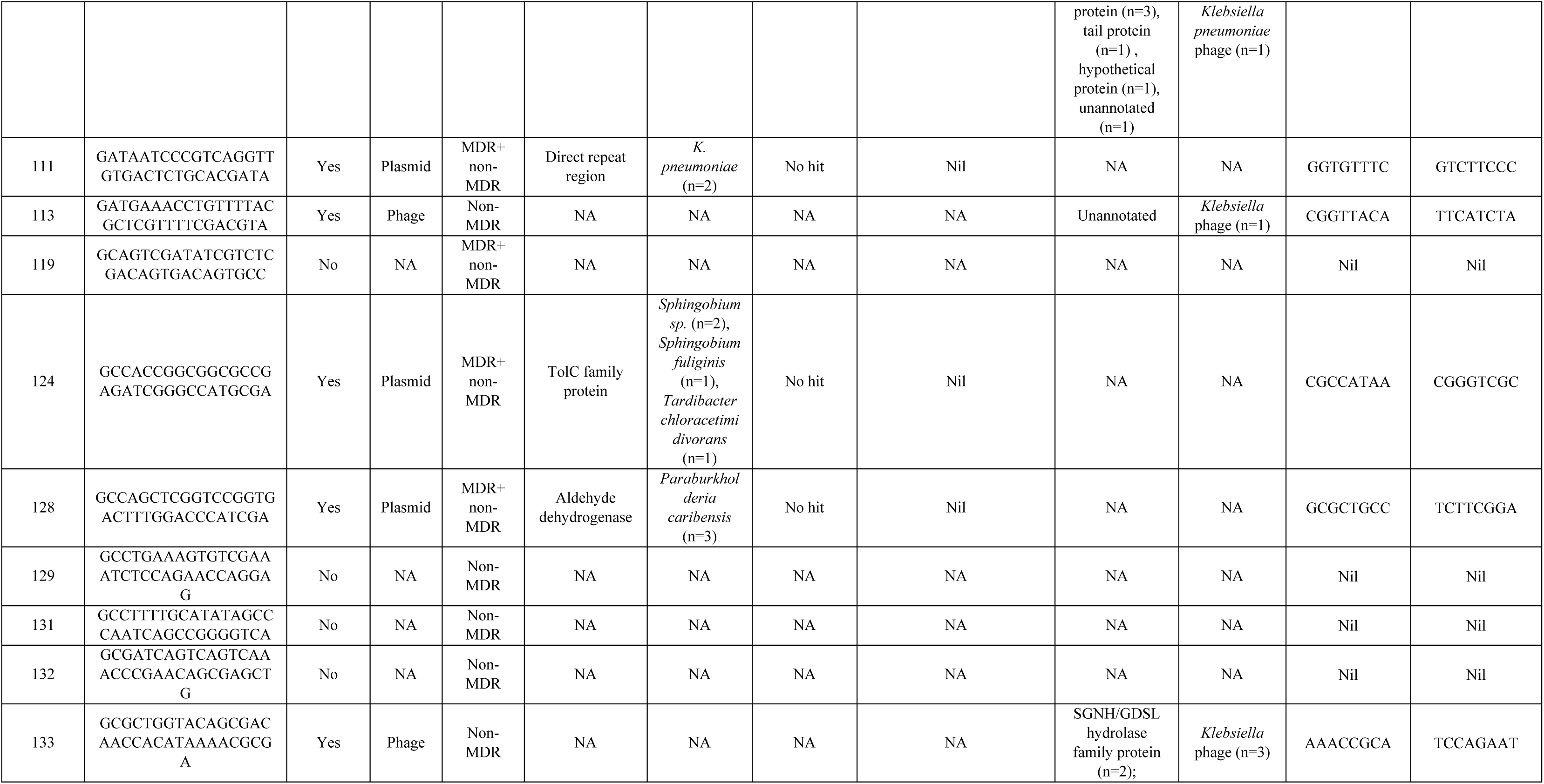
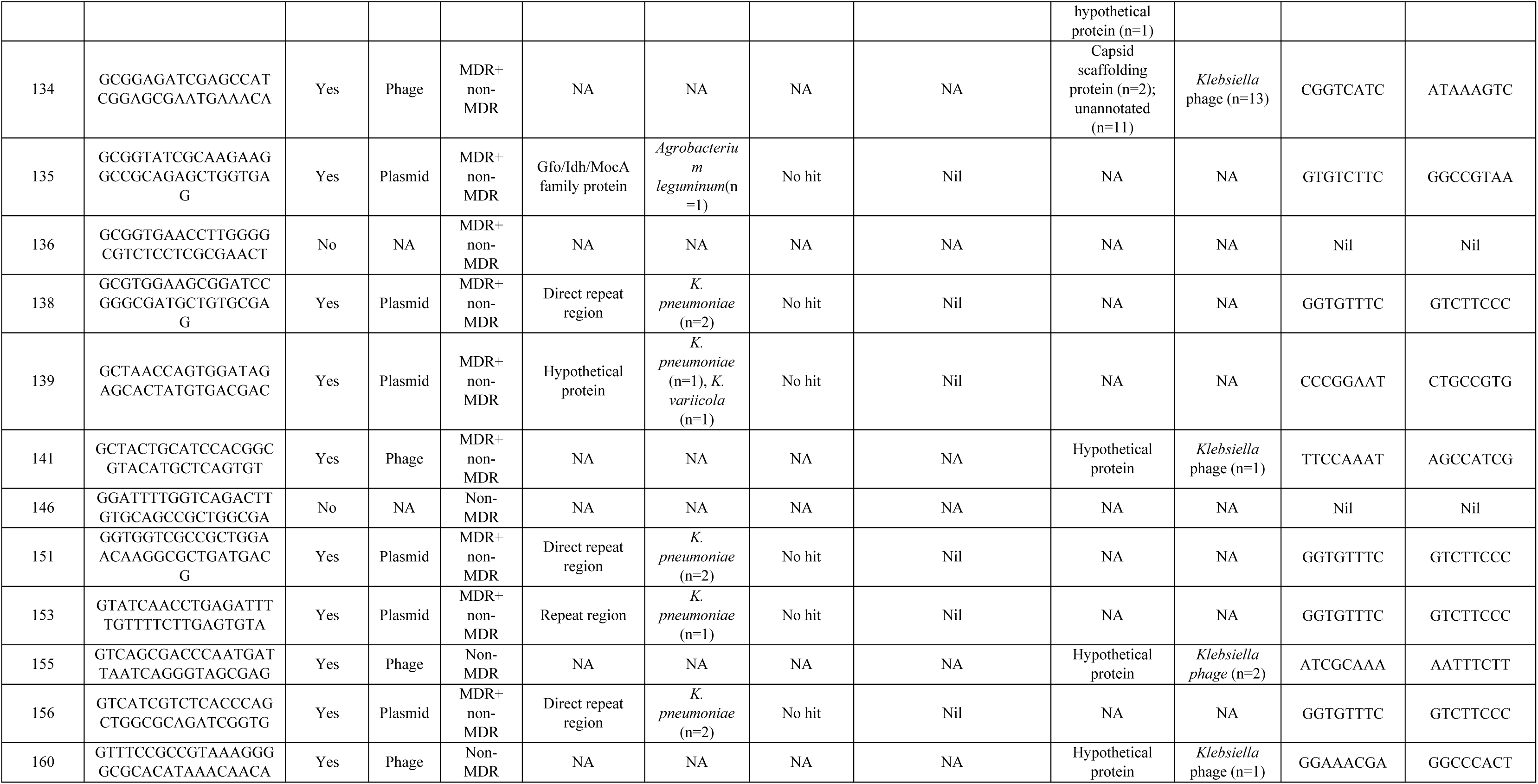
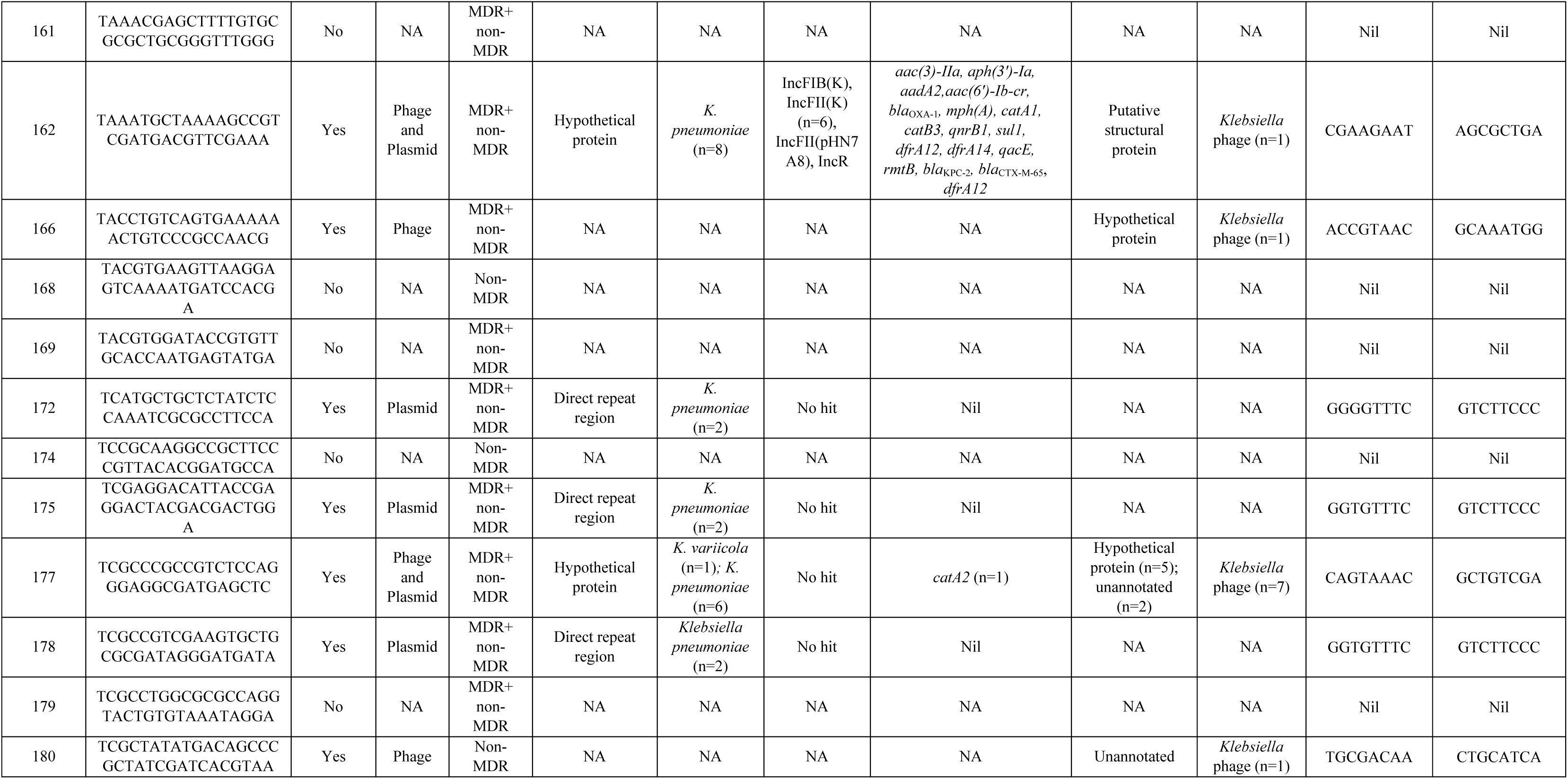
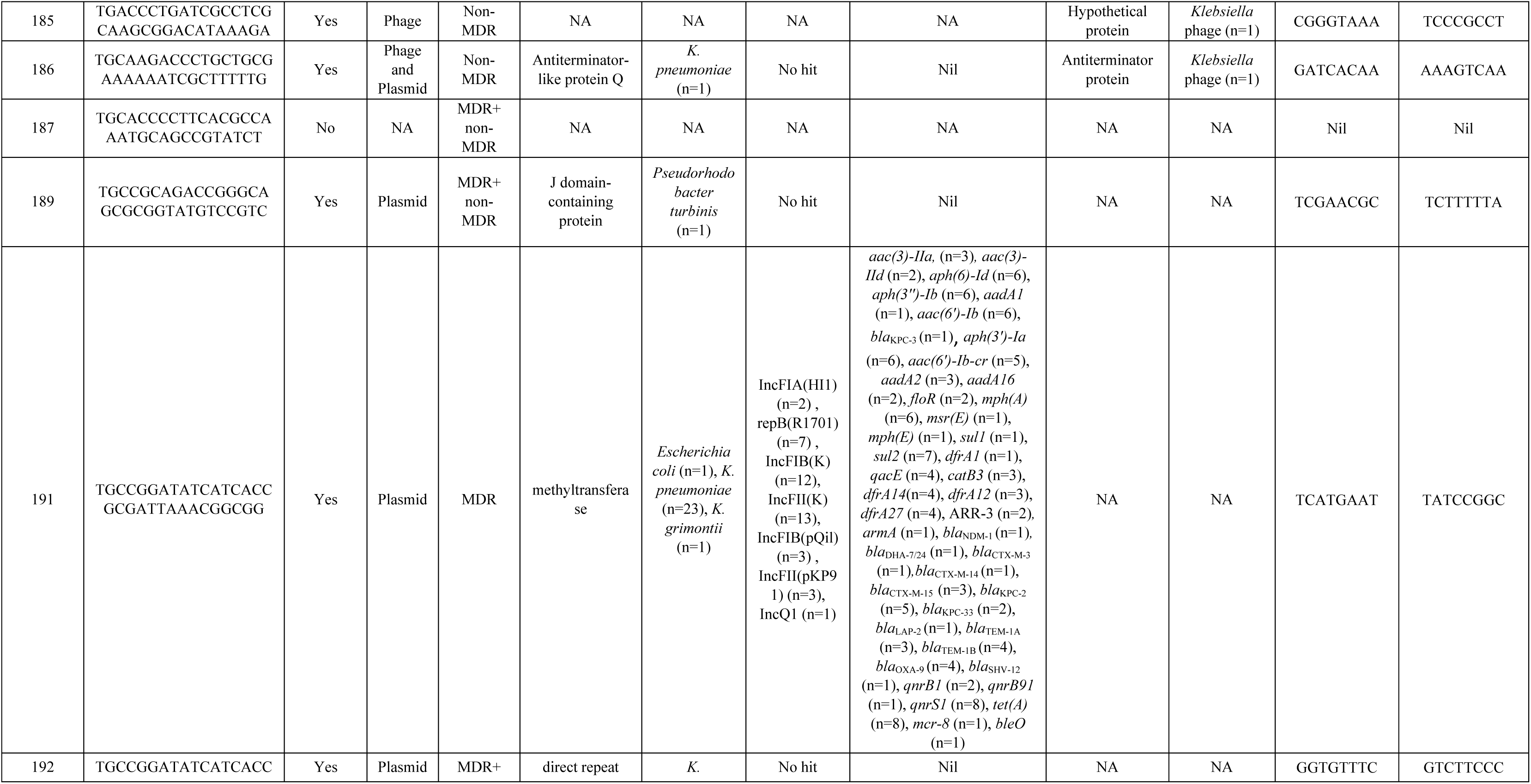
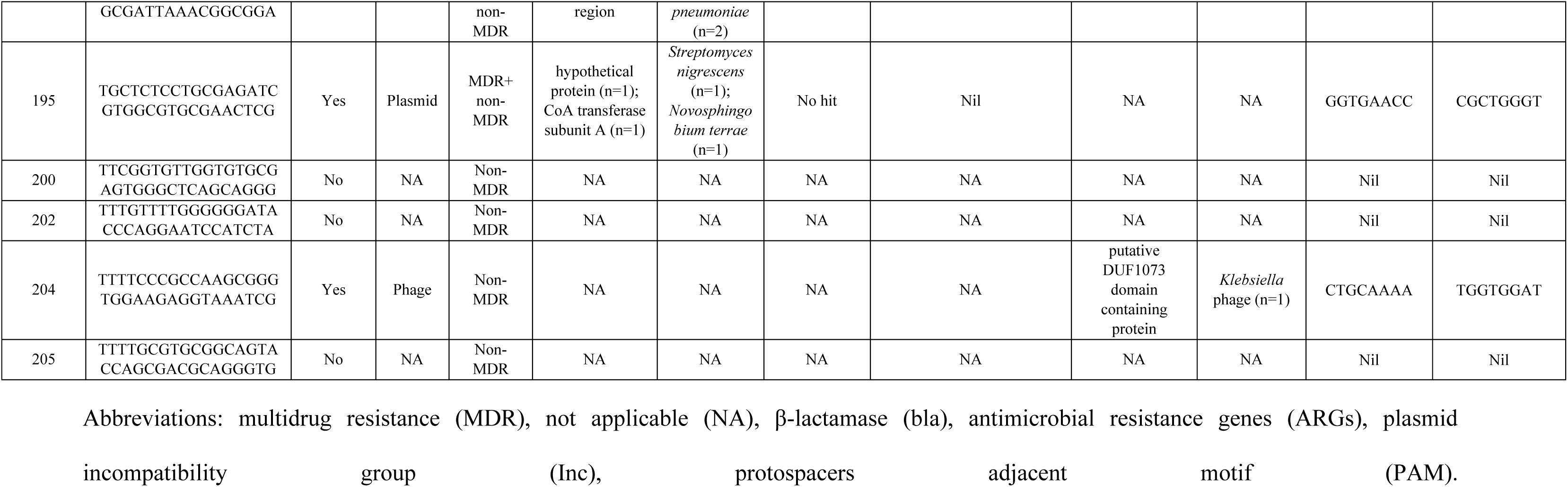
Characterization of distinct spacers (n=205) comprising of (A) unique spacers and (B) non-unique spacers identified from genome sequenced study *K. pneumoniae* (n=28) in terms of targeted protospacers, protospacer adjacent motifs (PAM), plasmid and phage targeted, sources, resistance genes carriage and replicons

Only 28% (25/91) of these spacers targeted plasmids which bore ARGs for β-lactams (*bla*_CTX-M,_ *bla*_SHV_), carbapenems (*bla*_KPC,_ *bla*_NDM_), chloramphenicols (*catA*), aminoglycosides (*rmtB, aac-6’-Ib-cr*), colistin (*mcr-8, mcr-10*), etc. Out of 25 ARG-bearing plasmid-targeting spacers, 11 targeted *bla*_KPC_-bearing plasmids and two targeted *bla*_NDM_-bearing plasmids. Certain spacers (spacer nos. 17, 34, 45, 54, 149, 165, 177) targeted plasmids which had single ARG (*bla*_SHV_/*catA2*), while spacer no. 191 targeted 44 ARGs of different categories including carbapenemase genes: *bla*_NDM_ and *bla*_KPC_, present on several plasmids (Figure 2, Tables 4.A-B and S5). Spacer nos. 62 and 191, retrieved from *bla*_NDM_-harboring study strains, could target protospacers present on other *bla*_NDM_-bearing plasmids. To understand why these two strains harbored spacers which targeted protospacers of other NDM-bearing plasmids but still could retain *bla*_NDM_ plasmids themselves, the protospacer sequences were analyzed. This revealed that in these two cases, the protospacers that have been targeted were absent in the *bla*_NDM_ plasmids present in the study strains, implying non-self-targeting nature of these spacers.

Seventeen plasmid-targeting spacers (19%, 17/91) targeted plasmids belonging to 17 different replicon types (Table S5). IncR plasmids were targeted by most of these spacers (53%, 9/17), followed by IncFII(pHN7A8) (47%, 8/17), IncFII(K) (47%, 8/17) and IncFIB(K) (41%, 7/17) (Tables 4.A-B, S5). Spacers could target protospacers in multiple replicon *viz*. spacer no. 191 could target seven different replicons (IncFIA(HI1), repB(R1701), IncFIB(K), IncFII(K), IncFIB(pQil), IncFII(pKP91), IncQ1) (Tables 4.B and S5).

### 3.11. Detailed analysis of phage-targeting spacers

Within the entire collection of spacers, phage-targeting spacers (n=64) i.e. spacers targeting only phages (n=42), as well as those targeting both phages and plasmids (n=22) were also noted. Out of all the phage-targeting spacers, 91% (n=58) were found to target phages specific for only 8 different bacteria (*Citrobacter sp., Enterobacter sp., Escherichia sp., Klebsiella sp., Proteus sp., Pseudomonas sp., Raoultella sp., Salmonella sp.*) where most of the targeted phages were *Klebsiella* sp. phages. Most of the phage-targeting spacers were found to primarily target hypothetical proteins (44%, 28/64) followed by non-coding regions (6%, 4/64). Additionally, certain genes encoding phage-specific proteins were also targeted along with protospacers coding for capsid scaffolding protein, tail length tape-measure protein H, putative tail assembly protein etc. (Tables 4.A-B).

Contrary to the plasmid-targeting spacers which targeted a wide range of bacteria (n=43), phage-targeting spacers could target phages specific for only 8 different bacteria (Tables 4.A-B). Hence, plasmid-targeting spacers possessed a greater target diversity compared to phage-targeting spacers.

### 3.12. Deciphering PAM specificity in complete type I CRISPR-Cas possessing study and global strain

PAMs flanking the 5ʹ and 3ʹ regions of the protospacer, function as markers for spacer adaptation and targeting. Assessment of PAMs were carried out in both local and global CRISPR-positive strains. In the local study strains, 229 diverse PAM sequences were noted, i.e., 113 and 116 PAMs at the 5ʹ- and 3ʹ-end of the protospacers, respectively (Tables 3, 4.A-B, S4). 5ʹ-GGTGTTTC-3ʹ (10%, 22/229) was the most frequently occurring 5ʹ-end PAM sequence and in case of 3ʹ-end PAM, 5ʹ-GTCTTCCC-3ʹ (8%, 18/229) was most frequent (Table S4). A difference in average number of identified PAMs were noted in non-MDR: 17.9 (179/10) v/s MDR: 5.8 (106/18) (Table 3), indicating increased identification of PAMs and hence, enhanced targeting by spacers of non-MDRs compared to MDRs (Tables 3, S4). PAMs were mostly found on plasmids and were frequently identified by spacers from non-MDRs (Table S4).

Out of 358 global strains with spacer arrays, 105 different PAM sequences could be identified and were located in both 5ʹ- (n=56) and 3ʹ-ends (n=49) of the protospacers. 5ʹ-GAAACACC-3ʹ, 5ʹ-GTTCGGCG-3ʹ and 5ʹ-ATTCGTTT-3ʹ were most frequent PAMs (n=27), followed by 5ʹ-ATTCGTTT-3ʹ (n=15) while 5ʹ- CCGTGCGC-3ʹ was most frequent (91%, 96/105), followed by 5ʹ-AGGCATAC- 3ʹ & 5ʹ-GTCTTCCC-3ʹ (77%, 81/105), and 5ʹ-GTCTTCAC-3ʹ (66%, 69/105) towards the 3ʹ-end, in the entire collection. Diversity of PAM sequences were high in the local strains compared to the global strains.

### 3.13. Phylogenetic analysis of *cas* genes among study and global strains

Phylogenetic trees for different *cas* genes i.e., *cas1, cas2, cas3, cas6, cse1* and *cse2* of all the global complete CRISPR-Cas containing *K. pneumoniae* strains and genome-sequenced study strains were constructed. For making the phylogenetic trees, two approaches were undertaken. Firstly, phylogenetic trees were made using the entire collection of global and local strains (data not shown). Strains in each tree clubbed into 4-6 clusters. Distinct global strains from each cluster and all study strains were selected and new phylogenetic trees were made (Figures 4.A,B). This was done for better interpretation of the phylogenetic relationship among the subtypes of type I CRISPR-Cas system in terms of the *cas* genes. From the different phylogenetic trees, it was observed that the *cas* genes were distinct among the subtypes (I-E and I-E*) but within their respective trees, they exhibited sequence similarities and formed clusters (Figures 4.A,B). Type I-E* strains formed 2 major clusters among all the *cas* genes whereas type I-E strains formed 3 major clusters in all trees except *cas6*, which contained 4 type I-E clusters. Type I-E* study strains (MDR and non-MDR) and few global strains formed one major clade, clade-1 (Tables 4.A, B). Type I-E strains-bc3c964 and SB615 formed a separate clade in all of the phylogenetic trees, showing major sequence variations within *cas* genes compared to other type I-E strains. Globally reported strains exhibited divergent clustering patterns, both inter- and intra-clade i.e., strains 04A025 (for *cas1, cas2, cas3, cas6, cse1* and *cse2*), 2018S07-013 (for *cas1* and *cas2*), ALRG-4569 (for *cas2* and *cas6*), WRC28_CMC387MPCH (*cas6* and *cse1*) and ATCC 43816 (for *cas2* and *cse2*) showed intra-cluster divergence, being placed at different locations of the clade-1 (type I-E*) for different *cas* genes (Figure 4.A, B). Similarly, type I-E* strains-DT12 and E16KP0035 broke clustering norms for *cse1* gene, clustering in the clade-3 with type I-E strains NKP1, DT1, 002SK2 and C51 (Figure 4B).

**Figure 4:**
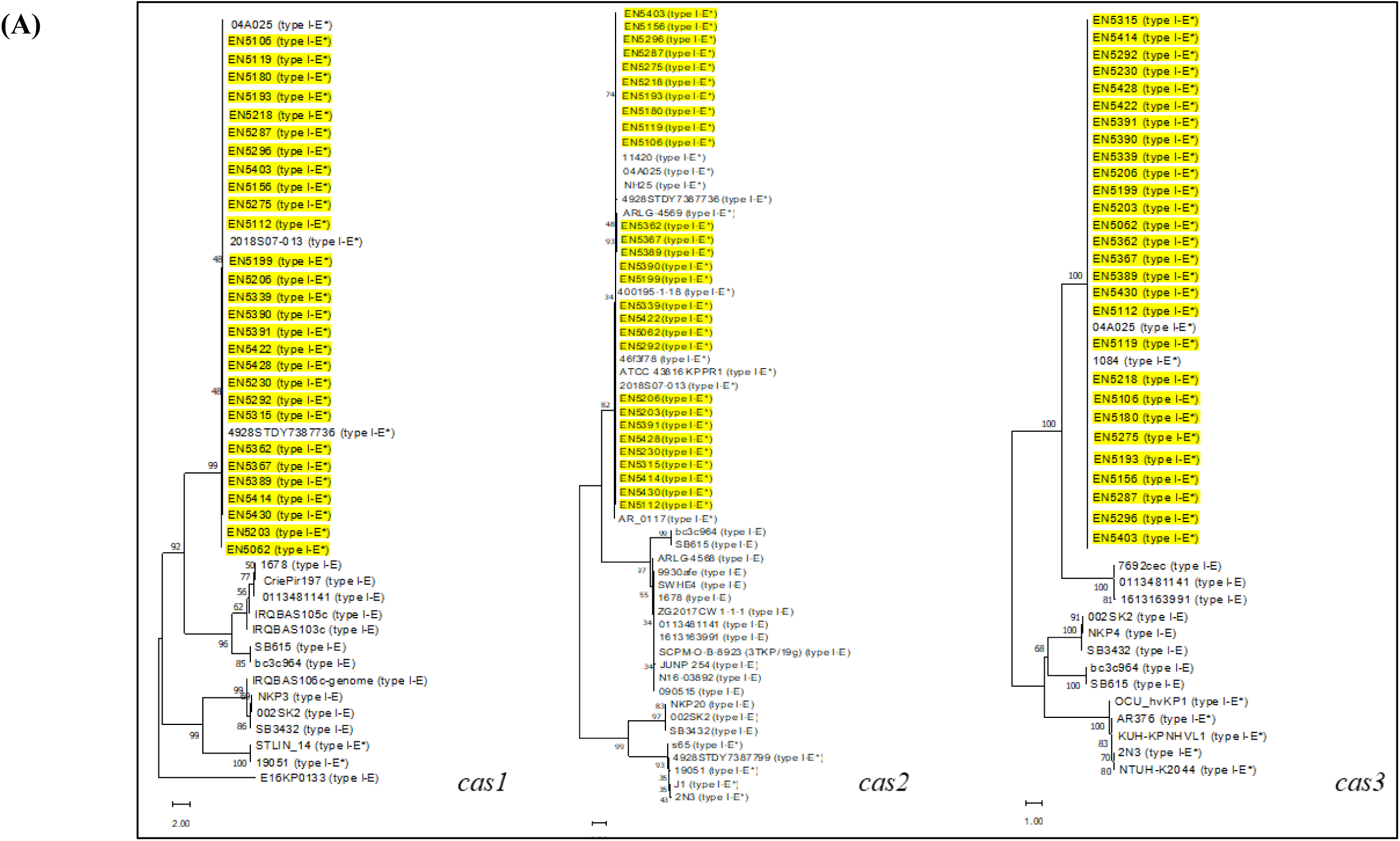
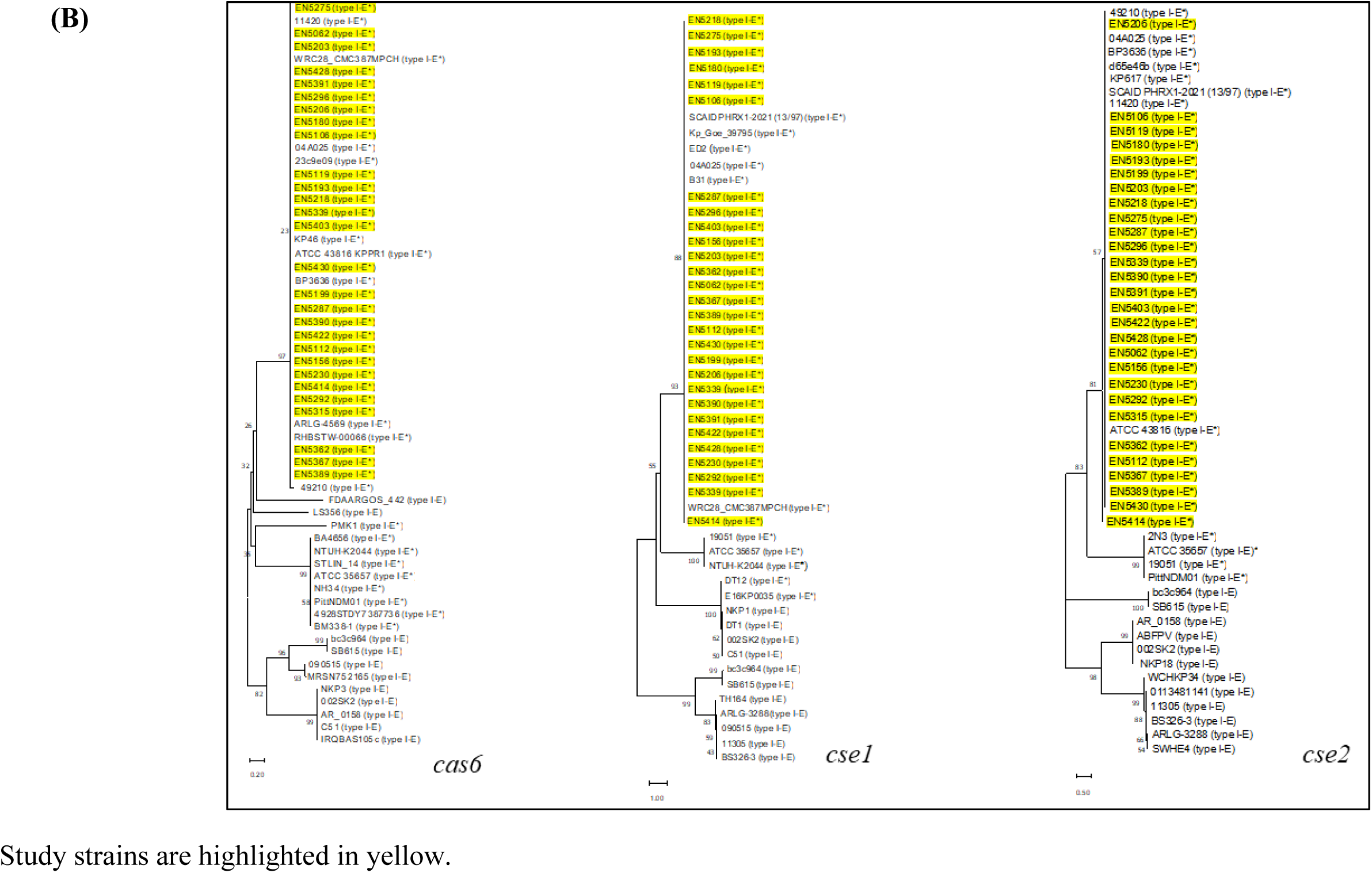
Phylogenetic comparison of different *cas* genes (*cas1, cas2, cas3, cas6, cse1* and *cse2*) among study and global CRISPR-Cas-positive *K. pneumoniae*.

## Discussion

CRISPR-Cas systems function as bacterial defense mechanisms that limit the acquisition of foreign DNA, such as from bacteriophages and plasmids (7). Given that ARGs are often spread via horizontal gene transfer, several studies have investigated the potential role of CRISPR-Cas in restricting their acquisition (13–16). Understanding this relationship could inform strategies to curb the spread of ARGs. While most studies (14–19) report an inverse relationship between CRISPR-Cas presence and MDR phenotypes i.e., absence of CRISPR-Cas in MDR strains, a few (20, 21) have observed the opposite trend, i.e., presence of CRISPR-Cas in drug-resistant strains. However, significant gaps remain in our understanding of CRISPR-Cas systems’ in MDR and non-MDR bacteria. Many studies have been limited to binary analyses of CRISPR-Cas and ARG presence or absence (14–17, 33), without delving into spacer diversity, target specificity (e.g., plasmids vs. phages), or associations with specific bacterial lineages. This study addressed these gaps by systematically analyzing lineage specificity of CRISPR subtypes, spacer diversity, protospacer origins (plasmid or phage), and ARG carriage within targeted plasmids across MDR and non-MDR strains.

The presence of complete type I CRISPR-Cas system in local study strains (25%) and the global strains analysed (22%) were consistent with a meta-analysis involving 1607 *K. pneumoniae* genome sequences (25%) (34) and previous reports from China 12.4% to 30.7% (15, 17, 33, 35, 36). Most studies have analysed *K. pneumoniae* of nosocomial origin between a period of 4 months to 6 years (15, 17, 33, 35, 36) whereas this study evaluated strains collected over a period of 13-years, providing a longitudinal perspective on CRISPR persistence. Prevalence of CRISPR-Cas system across this timeframe has remained relatively constant i.e., during two block years of 2008-2014 (24%) and 2015-2020 (26%), the prevalence was comparable.

Two subtypes of type I CRISPR-Cas (I-E and I-E*) have been reported in *K. pneumoniae* (34) which was also validated from our analysis of the global dataset. In the global dataset, dominance of type I-E* (67%, 129/192) in Asia and type I-E (69%, 42/61) in Europe respectively were noted. Similarly, phylogenetic comparison of *cas* genes among the strains from this study and global dataset also showed clustering of type I-E* strains together. Type I-E* was also dominant in strains from bloodstream infections, suggesting subtype adaptation to invasive niches or clonal lineage association. While among other global strains of non-blood (clinical) and environmental or zoological origin, dominance of both subtypes was noted. The data of the local strains are corroborated by global trends, as the dominance of the type I-E* subtype in strains from India and bloodstream infections aligns with findings from the global dataset.

The type I-E* CRISPR-Cas systems identified in the local study strains were found across 10 distinct STs, most of which belonged to epidemic high-risk AMR (ST14, ST15) and hypervirulent (ST23, ST2096) clones, mirroring the distribution observed in global type I-E* strains. This observation aligns with previous studies showing presence of type I-E* CRISPR in ST14, ST15, and ST23 (17, 36, 37). Global dataset from this study showed predominance of ST147 among type I-E, as also noted in a previous study (38). This study, similar to few other studies (17, 36, 37, 39), reports a potential linkage between specific STs and CRISPR-Cas subtypes in MDR strains.

This study did not find any statistically significant association between the presence of CRISPR-Cas systems and MDR phenotypes, which contrasts with several earlier reports (14–19). In globally disseminated MDR clones such as ST11 and ST258, absence of CRISPR was documented (18, 37). However, consistent with our results, findings by Gophna *et al.* supported the idea that CRISPR-Cas may persist and function alongside AMR determinants, particularly when under strong antibiotic selection pressures (40). When delving deeper beyond a simple presence/absence assessment of CRISPR-Cas and antibiotic resistance, several notable patterns emerged. Non-MDR strains exhibited higher spacer range (20–48 compared to 15–30 in MDR strains) and showed diversity in spacers (non-MDR:103 v/s MDR:38) as well as in their PAM sequences (non-MDR:179 v/s MDR:106). Even though the expression of *cas1* and *cas3* genes were comparable in MDR and non-MDR, the abovementioned features of non-MDRs collectively suggest the role of CRISPR-based interference and, consequently, a more limited acquisition of ARGs in non-MDR strains.

It should be noted, however, that the difference in spacer range between non-MDR and MDR strains was not observed in the type I-E* global dataset. This may be because, for local isolates, unique non-clonal strains were specifically chosen, whereas this criterion could not be applied to the global dataset, and hence a possible skew in the global results cannot be excluded. Additionally, targets of CRISPR spacers were more variable in non-MDR strains. Many of these spacers targeted Enterobacterales and promiscuous plasmids-potent vehicles for the horizontal spread of ARGs- and most such targets were recognized by non-MDR strains. This, together with the observed diversity in PAM sequences, further supports the notion that CRISPR-Cas systems may be contributing in restricting the acquisition of ARGs in non-MDR strains. Notably, many of the spacers that identified PAM sequences were directed against plasmids known to carry ARGs, underlining their potential to enable effective recognition and removal of invading genetic elements.

Our findings underscore that the presence of CRISPR-Cas systems alone cannot account for ARGs present, which may explain the inconsistencies observed across earlier studies. Higher average spacer diversity and a greater number of spacers targeting ARG-bearing plasmids in non-MDR study strains is indicative of their role in reduced ARG content in the non-MDR strains but it is not conclusive. Despite possessing CRISPR-Cas systems, our MDR strains had acquired diverse ARGs not represented in their spacers, suggesting that in environments with a high ARG reservoir- such as hospitals- the likelihood of acquiring untargeted ARGs is significantly increased. This highlights the role of environmental exposure in shaping both spacer acquisition and subsequent excision of foreign genes, with additional influences likely from restriction-modification systems and anti-CRISPR mechanisms. The extended 13-year sampling of *K. pneumoniae* strains, although from a single centre, provides a longitudinal perspective that complements and, in some respects, exceeds the temporal scope of prior nosocomial studies. This also indicates that prevalence of CRISPR has remained to about 25% across these many years. Together, these observations not only enhance our understanding of CRISPR-AMR dynamics, particularly of the spacer and target diversity but also suggest that designing CRISPR-Cas tools to target priority ARGs could be a viable approach to limit their spread.

## Acknowledgement

We thank Dr. Martin Rethoret-Pasty, curator for the database hosted on Institut Pasteur (https://bigsdb.web.pasteur.fr) for curating the *Klebsiella pneumoniae* sequence type and making it publicly available. We extend our thanks to Dr. Surajit Basak, for his scientific advice. We also thank the staff of the Department of Neonatology, who cared for the neonates, Subhadeep De and Saikat Sahoo for their laboratory assistance and data maintenance.

## Data availability

Whole genomes sequences of the study isolates have been submitted to NCBI database under BioProject number: PRJNA548120

## Transparency declarations

None to declare.

